# Integrated analysis of COVID-19 multi-omics data for eQTLs reveals genetic mechanisms underlying disease severity

**DOI:** 10.1101/2024.12.18.629144

**Authors:** Jeongha Lee, Eun Young Jeon, Liyang Yu, Giovanna Durante, Hye-Yeong Jo, Sang Cheol Kim, Woong-Yang Park, Hyun-Young Park, Siming Zhao, Murim Choi

## Abstract

**Background:** The global pandemic caused by the SARS-CoV-2 virus provided an unprecedented opportunity to investigate genetic factors influencing the disease severity of the viral infection. Despite a plethora of recent research on both SARS-CoV-2 and COVID-19, few have taken a systems biology approach to address individual-level variation, especially based on non-European populations.

**Methods:** We analyzed multi-omics data generated at three timepoints from 184 Korean COVID-19 patients with mild or severe symptoms, composed of whole genome sequencing, blood-based single-cell RNA-sequencing (2.15M cells), 195 cytokine profiles, and human leukocyte antigen (HLA) allele data. Expression quantitative trait loci (eQTLs) were called and their biological implications were evaluated.

**Results:** We identified disease severity interacting eQTLs (*n* = 404) and disease progression interacting eQTLs (*n* = 972) for various cell types. We elucidated a complex regulatory mechanism involving *HLA* genes and their targets, and identified genetic determinants of cytokine levels. Finally, we show how regulation of interaction-eQTLs (ieQTLs) is established by upstream transcription factors (TFs), illustrating complex regulation of the *IGFBP7* ieQTL, which is potentially important in conferring differential severity. Our COVID-eQTLs were independently validated against a large-scale single-cell PBMC eQTL dataset (1M-scBloodNL), providing robust cross-dataset replication across cell types.

**Conclusions:** This study illuminates an efficient molecular interrogation framework that can be applied toward understanding infectious disease progression in individuals of different genotypes.

## Background

Despite their importance in human biology and diseases, elucidation of the genetic mechanisms that influences individual variation in susceptibility to infection and the severity of infectious diseases has been challenging due to the high contribution of non-genetic factors (*i.e.*, low heritability) and involvement of multiple genetic variants [1]. The emergence of severe acute respiratory syndrome coronavirus 2 (SARS-CoV-2) and associated global Coronavirus disease 2019 (COVID-19) pandemic offers an unprecedented opportunity to recruit and study large cohorts across diverse ethnic groups and conduct a genetic evaluation of virus-driven disease susceptibility and severity.

Several factors, including age, sex, and comorbidity are known to contribute to COVID-19 susceptibility and severity [2–5]. To address the genetic contribution of differential severity in addition to these factors, the COVID-19 Host Genetics Initiative (HGI; https://www.covid19hg.org/) determined genome-wide associated variants from more than 220K infected individuals with differing COVID-19 severity [6]. Another effort took a rare variant analysis approach and elucidated a list of genes for which certain alleles confer severe symptoms upon the infection [7,8]. However, neither of these studies examined the complex networks of genomic variants and gene expression that are cell-type-specific, which may eventually affect disease severity and prognosis. Although recent international meta-analyses have included non-European cohorts, ancestry representation remains uneven. Indeed, COVID-19 studies using non-European ancestries have successfully identified novel signals that play roles in the virus infection and disease progression [9,10].

Analysis of expression quantitative traits (eQTLs) allows for identification of genetic variants that influence gene expression in a tissue [11–14], which can reveal novel insights into the genetic regulation of gene expression and explain disease-associated GWAS loci (*i.e.*, colocalization) [15–18]. Early COVID-19 colocalization studies largely relied on eQTL reference panels derived from healthy tissue or *ex-vivo* stimulated PBMC dataset and therefore could not fully capture infection-induced regulatory effects [19–21]. More recent patient-based bulk and single-cell studies have begun to identify disease-state-dependent eQTLs that colocalize with COVID-19 GWAS signals [22,23]. Also, a recent study provided genetic evidence that not all GWAS loci can be tied to an eQTL as GWAS loci and eQTLs are inherently different in many genetic aspects [24], which may explain the low colocalization rate (20-30%). Moreover, most of the previous eQTL studies lacked clinical implication due to analyses being performed on healthy tissues, or on a combined set of healthy and diseased tissues. To circumvent this limitation, we and others have extended the eQTL approach to identify eQTLs that are functional in a disease-, condition-, or cell-state-specific manner. The resulting response-eQTLs and interaction-eQTLs (ieQTLs) have demonstrated their utility in elucidating novel, individual-specific genetic pathways and drug targets [12,13,25]. Furthermore, our multi-omics approach allows the probing of genetic mechanisms underlying the expression of *HLA* genes and cytokines, which are critical factors in the determination of disease severity.

In this study, we analyzed multi-omics data from 184 Korean COVID-19 patients. The patients were classified by disease severity into the mild and severe groups, and serial sampling and data generation yielded 665 runs of single-cell RNA sequencing (scRNA-seq) and 501 runs of cytokine levels. Both eQTLs and ieQTLs, collectively termed as COVID-eQTLs, were called and integrated with genetic loci previously associated with COVID-19. Through this, we uncovered a number of molecular mechanisms explaining the association of previously-identified or novel factors with infection status. Our approach further brought to light an ieQTL that regulates *IGFBP7* expression through the transcription factor (TF), PATZ1, which may contribute to differential severity of disease. Overall, these variants may inform the prediction of the severity of virus infection.

## Methods

### Cohort and data generation

Details of patient recruitment, patient classification, sample collection, clinical and genetic data generation and processing procedures were described previously [26]. The study protocol was approved by the Institutional Review Boards (CNUH 2020-12-002-008, SEOUL 2021-02-016, SMC-2021-03-160, and 2020-09-03-C-A). All the participants provided written informed consent [26].

Most patients were infected with the B.1.497, B.1.619, or B.1.620 strains, reflecting the major COVID-19 lineages at the time of the patient recruitment [26]. Classification of patients by COVID-19 severity was performed following the NIH treatment guidelines [27]. Patients classified as “severe” presented COVID-19 symptoms with pneumonia and one of the following: respiratory rate of more than 30 breaths/min, oxygen saturation <93% at room temperature, <300 mmHg of PaO2/FiO2, or rapid progression of infiltration (>50%) observed by chest computed tomography imaging. In this study, the Mild group is composed of asymptomatic (*n* = 11), mild (*n* = 34), and moderate (*n* = 47) cases, and the Severe group of severe (*n* = 55) and critical (*n* = 37) cases, as defined previously (Supplementary Table 1) [26].

Use of the data in this study was approved, and the data downloaded from the Clinical & Omics Data Archive (CODA), Korea National Institute of Health, Republic of Korea (https://coda.nih.go.kr/), under the title of “Establishment of multi-omics data related with COVID19”.

### scRNA-seq data processing

PBMCs were acquired to construct scRNA-seq libraries using the 10X Chromium Next GEM Single Cell 5’ Library kit. Generation of scRNA-seq matrix was described in a previous study [26]. PBMCs from four participants were pooled together at the equal number of cells. Doublet detection was performed using Scrublet to identify potential doublets in the data [28]. Single-cell RNA-sequencing data from patients were processed using Scanpy v1.6.0 [29] and Seurat v5 [30]. Cells that expressed <200 genes were excluded from the analysis, and genes expressed in fewer than three cells were removed. Cells with >15% mitochondrial RNA content were likewise excluded from further analysis. Data normalization, variable feature selection, and dimensionality reduction were performed with batch effect correction using Harmony [31]. Clustering was conducted at a resolution of two, and uniform manifold approximation and projection (UMAP) was used for visualization. Cell type annotations were obtained via Azimuth [32], and the sketched assay was projected onto the full dataset for comprehensive integration.

### WGS data processing

To call genetic variants in the participants, WGS reads were aligned to the human genome reference build GRCh38. gVCF files were generated as described previously[26] and subsequent steps for consolidation of gVCFs, joint-calling, and variants recalibration were conducted using the GATK 4.1.6.0 “germline short variant discovery” pipeline [33]. Only variants passing the recalibration step were included. Additional filtering steps to remove low-quality variants included, excluding low-complexity regions [34], applying allelic balance cutoffs of 0.22 for SNPs and 0.2 for indels, restricting indel sizes to ≤50bp, and including only biallelic variants. Samples were pruned based on sex discordancy, missing genotype rate >0.05, kinship coefficients >0.177, and population stratification adjustments using PCA (Supplementary Figure 1). Population stratification was evaluated using the 1000 Genomes Project and the Korean Variant Archive (KOVA) [35].

### HLA allele type processing and association analysis

HLA genotyping was performed as described previously [26]. HLA alleles were analyzed at two-field resolution and encoded as allele dosages. Alleles with ambiguous or missing HLA calls were excluded, and only alleles carried by at least five individuals were tested. Associations between HLA allele type and infection status were assessed using multivariable logistic regression adjusted for age, sex, and the first genotype-derived principal components. Odds ratios and 95% confidence intervals were calculated from regression coefficients.

### Pseudobulk eQTL and interaction-eQTL calling

For eQTL analysis, samples with both genotype and phenotype data available were included. Preprocessing steps for eQTL input were conducted separately for each combination of condition and cell type. To ensure robust signal detection, seven major cell types were analyzed (B cells, CD4+ T cells, CD8+ T cells, NK cells, dendritic cells, monocytes, and other T cells). For each combination, count matrices were aggregated at the individual level, followed by quantile normalization across samples and inverse normal transformation for gene expression values. Corresponding genotype data for each combination were extracted and filtered based on the following criteria: missingness <5%, minor allele frequency (MAF) > 0.05, Hardy-Weinberg equilibrium *P* > 1.0 x 10^-5^, and LD clumping using an R^2^ threshold of 0.9, sorting variants by their MAF. Age, sex, the top three genotype PCs and expression PCs generated from PCAforQTL were used as covariates [36]. For the cis-eQTL mapping, TensorQTL was used with ‘cis_nominal’ mode [37].

To assess the sharing patterns of eQTLs across cell types and increase eQTL identification power, the Multivariate Adaptive Shrinkage (MASH) method was applied for each condition [38]. In this analysis, the 15,000 tests with the lowest ‘pval_nominal’ values from TensorQTL were designated as the strong set, and a random subset of 200,000 tests was used. Following MASH procedures, we learned correlation structure among null tests using the random subset and data-driven covariance matrices using the strong subset. We then fit the mashr model to the random subset to estimate prior parameters, and subsequently derived posterior summaries for the strong set and the complete dataset. eQTLs with local false sign rate (LFSR) under 0.05 were used for further analysis.

For the interaction-eQTL (ieQTL) analysis, either severity status (mild or severe) or timepoint (1,2, or 3) was used as the interaction term in a linear mixed model (LMM). The model was defined as follows:

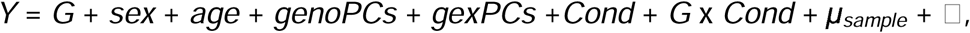

where Y refers to gene expression, G to genotype, and sex, age, genoPCs (genotype PCs), and gexPCs (expression PCs) are included as covariates. The term Cond refers to the condition variable (severity or timepoint), and μ*sample* represents the random effect for samples. For sv-ieQTL analysis, one LMM with severity as the condition variable was constructed (M-S). To capture additional ieQTL signals, pairwise comparison of severity-specific conditions (M1-S1, M2-S2, and M3-S3) was also performed. For tp-ieQTL analysis, pairwise comparisons between timepoints (M1-M2, M1-M3, M2-M3, S1-S2, S1-S3, and S2-S3) were performed using the LMM model. To incorporate cell-type sharing patterns and increase power, MASH analysis was applied similarly as in eQTL calling, and a threshold of LFSR < 0.05 was used to determine statistical significance.

### Comparative analyses with other eQTL sets and GWAS

To assess the correlations of our eQTL data with external datasets, we obtained single-cell PBMC eQTL data from the 1M-scBloodNL dataset (Curion et al., Nat Genet 2024), as well as bulk whole blood eQTL data from the GTEx v8 dataset and the Japan Taskforce [23]. A lead eSNP per gene for each cell type and shared across M1, M3, S1, S2, and S3 was extracted based on nominal *P*-values from TensorQTL. Then, we searched for these lead eSNP-gene pairs in each external eQTL dataset and extracted the corresponding effect size and standard error. The proportion of true replicated associations (pi1 = 1 – pi0) was estimated by fitting an adaptive shrinkage model (ash function, ashr package) to these matched external effect sizes and standard errors, and alignment was quantified using Pearson correlation coefficients between internal and external effect sizes.

To evaluate colocalization of GWAS signal with our eQTL dataset, the COLOC package was used [39]. The priors for the probabilities of causal associations (p1, p2, and p3) were set to 1 x 10^-4^, 1 x 10^-4^, and 1 x 10^-5^, respectively. COVID-19 GWAS data were obtained from the COVID-19 Host Genetics Initiative (HGI; https://www.covid19hg.org/), while GWAS data for other immune diseases were sourced from the GWAS Catalog (https://www.ebi.ac.uk/gwas/ and Supplementary Table 2). Genomic regions with *P* ≤ 1.0 x 10^-5^ and extending 250 Kbp upstream and downstream of each locus were extracted. Genes within these regions were considered for analysis if they contained significant eQTL calls, as identified by MASH [38]. COLOC analysis was then conducted on these genes, provided there were at least 20 SNPs shared between the GWAS and eQTL datasets [40].

### Analysis of gene features

Gene Ontology, KEGG pathway, and gene set enrichment analysis (GSEA) analyses were performed for eGenes from each condition and cell type using clusterProfiler [41]. Gene members for Reactome pathways were downloaded from ‘https://www.gsea-msigdb.org/gsea/index.jsp’. Reactome pathway titles and IDs used in this study are as follows:

REACTOME_INNATE_IMMUNE_SYSTEM [R-HSA-168249; M1036]

REACTOME_ADAPTIVE_IMMUNE_SYSTEM [R-HSA-1280218; M1058]

REACTOME_CELL_CYCLE [R-HSA-1640170; M543]

REACTOME_MUSCLE_CONTRACTION [R-HSA-397014; M5485]

REACTOME_REPRODUCTION [R-HSA-1474165; M26956]

REACTOME_METABOLISM_OF_LIPIDS [R-HSA-556833; M27451]

To assess the enrichment of eQTLs in epigenomic markers and genomic annotations, we utilized the *fenrich* module from QTLtools [42]. Chromatin state data were obtained from the Roadmap Epigenomics Project using the 15-state model, focusing on the five immune cell types with cell annotations relevant to the project (*i.e.*, B [E032], NK [E046], Mono [E029], CD8+ T [E048, E047], and CD4+ T [E045, E044, E039, E037, E042]) [43].

Genomic annotations were performed according to the Comprehensive gene annotation from GENCODE release 26 [44]. All isoforms of each gene were collapsed into a single transcript using the *collapse_annotation.py* script from the gtex-pipeline provided by the Broad Institute (‘https://github.com/broadinstitute/gtex-pipeline/tree/master/gene_model’). Enrichment tests were performed on lead SNPs using QTLtools with the ‘--permute 1000’ option.

### Cytokine data processing, pQTL calling, and heritability estimation

Genotype data and quantifications of 195 cytokines data were collected from 395 individuals as described previously (Supplementary Table 3) [26]. Briefly, a customized panel based on the Luminex MAGPIX platform (LUMINEX Corporation, USA) was used to measure cytokines in plasma from the participants. For the Mild groups, data were available for 269 individuals at timepoint 1, 104 individuals at timepoint 2, and 187 individuals at timepoint 3. For the Severe group, data were collected from 126 individuals at timepoint 1, 113 individuals at timepoint 2, and 97 individuals at timepoint 3. Raw cytokine levels were first log-transformed and then mapped to genotype data using a linear regression model, with age and sex as covariates. Cytokines with zero values in more than 15% of the samples were excluded. To call protein QTLs (pQTLs), cytokine identifiers were converted to corresponding gene names, and the gene positions were annotated with GRCh38. Due to a high rate of false-positive calls, the region on chromosome 17 from positions 508,668 to 714,839 was excluded from the analysis. A Bonferroni-adjusted *P*-value of < 0.05 was used as the threshold to identify significant pQTLs.

Genomic heritability of cytokine levels was estimated using Genome-wide Complex Trait Analysis (GCTA) version 1.93.2 [45]. The genetic relationship matrix (GRM) was constructed for each cytokine using SNPs located within its *cis*-region, including only the Mild 1 and Severe 1 groups, by applying the --grm option. Log-transformed cytokine levels from all disease severities and timepoints served as the phenotypic variables. Phenotypic data for each cytokine were input separately in phenotype files. Age, sex, and severity were included as covariates for estimating the proportions of phenotypic variance explained by the SNPs. Heritability estimates were obtained using the restricted maximum likelihood method.

### Identification of upstream regulators of sv-ieQTLs

To identify putative upstream regulators of sv-ieQTL, firstly SCENIC[46] was used to identify regulon activities across cell types and conditions. Since there are 2.15M cells in our scRNA matrix, we leveraged the Python-based geosketch [47] to subset an informative set of 200K cells, retaining only cells with more than 200 genes, less than 4,000 genes, and less than 15% of transcripts of mitochondrial origin; additionally, only genes with expression detected in more than three cells were used. The sketched matrix was then extracted in loom format and utilized for SCENIC analysis.

Subsequently, we proposed several criteria to narrow the set of putative upstream regulators: (1) significant changes in TF DNA binding ability due to the presence of ieSNP, (2) absolute correlation more than 0.3 for regulon activity and ieGene expression across individuals, (3) absolute correlation more than 0.3 for regulon activity and eQTL effect size across the six conditions, and (4) regulon with activity biased to either mild or severe status. Binding motif occurrences were identified using FIMO from the MEME Suite program [48]. Flanking genomic sequences 15 bp upstream and downstream of each ieSNP were used as input, and binding was considered to occur for ieSNP-TF pairs with *P*-values less than 1 x 10^-4^. We used only those pairs that displayed a differential binding profile in relation to the ieSNP reference and alternative alleles. Regulon activity bias dependent on severity status was quantified by logistic regression using the *glm* function (R version 4.4.1). Degree of bias was estimated as the log of the odds ratio, with an absolute value of at least 50 considered to indicate differential regulon activity.

For visualization of epigenetic data in the UCSC genome browser, we downloaded call sets from the ENCODE portal[49] (https://www.encodeproject.org/) with the following identifiers: ENCFF060KZZ (ATAC-seq), ENCFF778SYZ (DNase-seq), ENCFF580XLE (CRE), ENCFF503MRH (H3K27ac ChIP-seq), ENCSR549PVK (PATZ1 ChIP-seq). ATAC-seq, DNase-seq, and H3K27ac ChIP-seq positions were converted to GRCh38 coordinates with ucsc-liftover.

## Results

### COVID-eQTLs signify COVID-19 severity

The collection of participants and generation of multi-omics data used in this study were previously described [26]. Briefly, Korean participants that were afflicted with COVID-19 during October 2020 to September 2022 were classified into the “Mild” (*n* = 92) or “Severe” (*n* = 92) groups based on the severity of host response to infection following the WHO guideline (Supplementary Table 1 and Supplementary Figure 1) [27]. Blood samples from each individual were collected three times at two-week intervals, with the first sampling performed on the day that the infection was confirmed; this resulted in six sample groups: mild timepoints M1, M2, and M3 and severe timepoints S1, S2, and S3. The omics data processed for this study included the following: blood-based scRNA-seq, whole genome sequencing (WGS), human leukocyte antigen (HLA) sequencing (HLA-seq), and blood-based cytokine profiles (Fig. 1A-B and Supplementary Table 4). Clustering and annotation of the scRNA-seq data, encompassing 2.15M cells from 665 samples across healthy controls and six disease points, revealed a slight increase in the proportion of monocytes (1.2-fold) and a decrease in CD4+ T cells (0.73-fold) in the Severe group compared to the Mild group (Fig. 1C; Supplementary Table 4), consistent with previous observations [50,51]. We also investigated the expression patterns of interferon-stimulated genes (ISGs) in each group across cell types using interferon gene sets from previous studies [52–54]. Consistent with those studies, ISG expression was elevated in the Mild group across all cell types, and the expression declined over time (Supplementary Figure 2). HLA allele analysis identified no statistically significant association with infection status (Fig. 1D). In contrast, principal component analysis (PCA) of circulating cytokine profiles revealed a severity-related gradient along PC1, with severe, mild, and uninfected control samples distributed sequentially (Fig. 1E). Using these data, we sought to understand the contributions of genetic variants in determining disease severity following SARS-CoV-2 infection through gene expression regulation. PCA-mediated clustering using effect size value of eQTLs from major cell type clusters revealed disease severity (represented by PC1 and PC2) to be a second determinator following cell type (represented by PC1) (Fig. 2A). Then we called ieQTLs which interact with COVID-19 infection status (inf-ieQTL), severity (sv-ieQTL), or progression time point (tp-ieQTL) (Fig. 2B-E). We will refer to all eQTLs and ieQTLs identified here as COVID-eQTLs.

**Figure 1.**
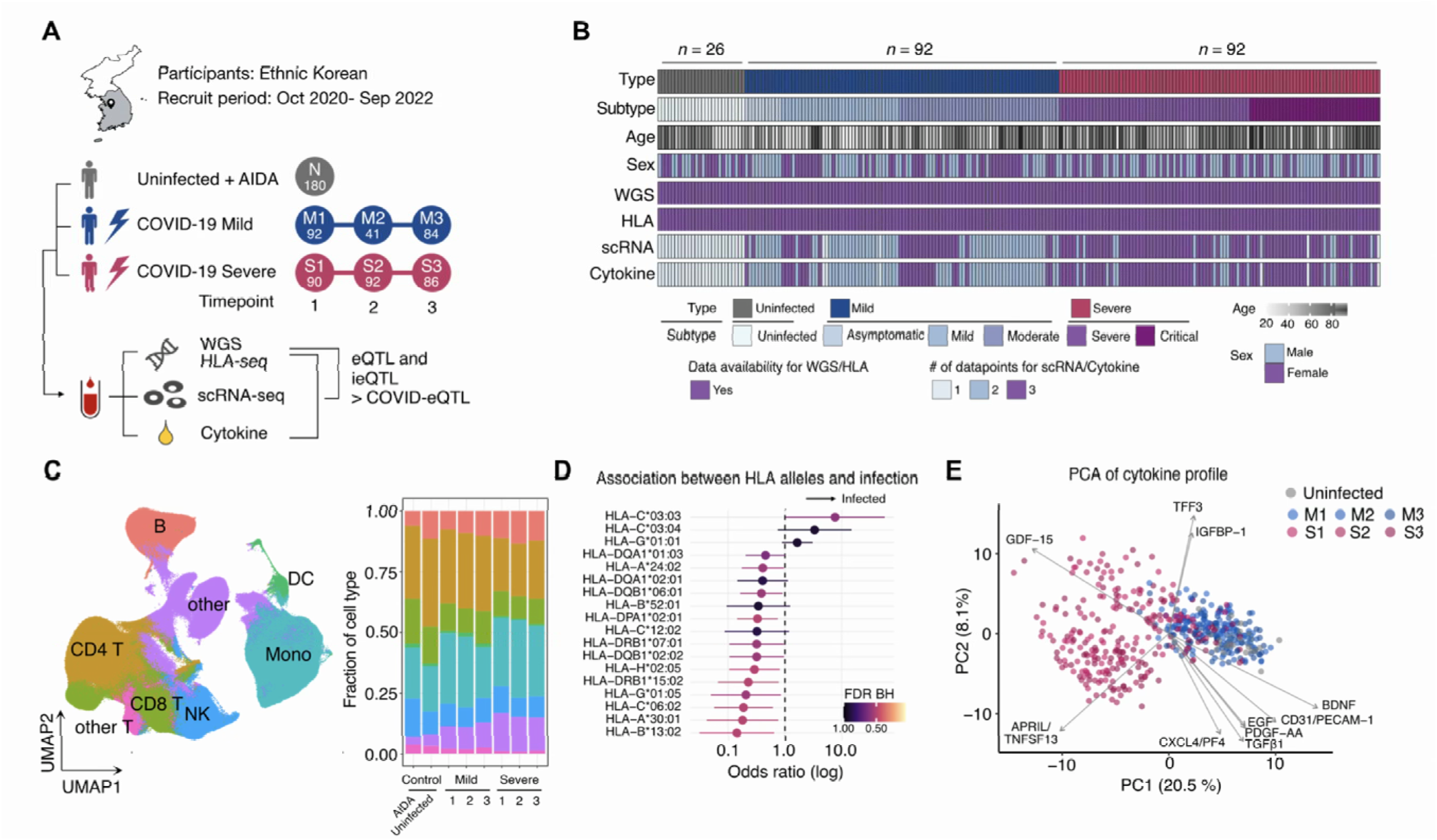
Study scheme and sample profile. (A) Cohort description and data generation scheme. (B) Data composition by sample. (C) Cell type composition UMAP based on PBMC scRNA-seq data. (D) Enrichment of HLA alleles between infected and non-infected individuals. (E) Serum cytokine profile by PCA.

**Figure 2.**
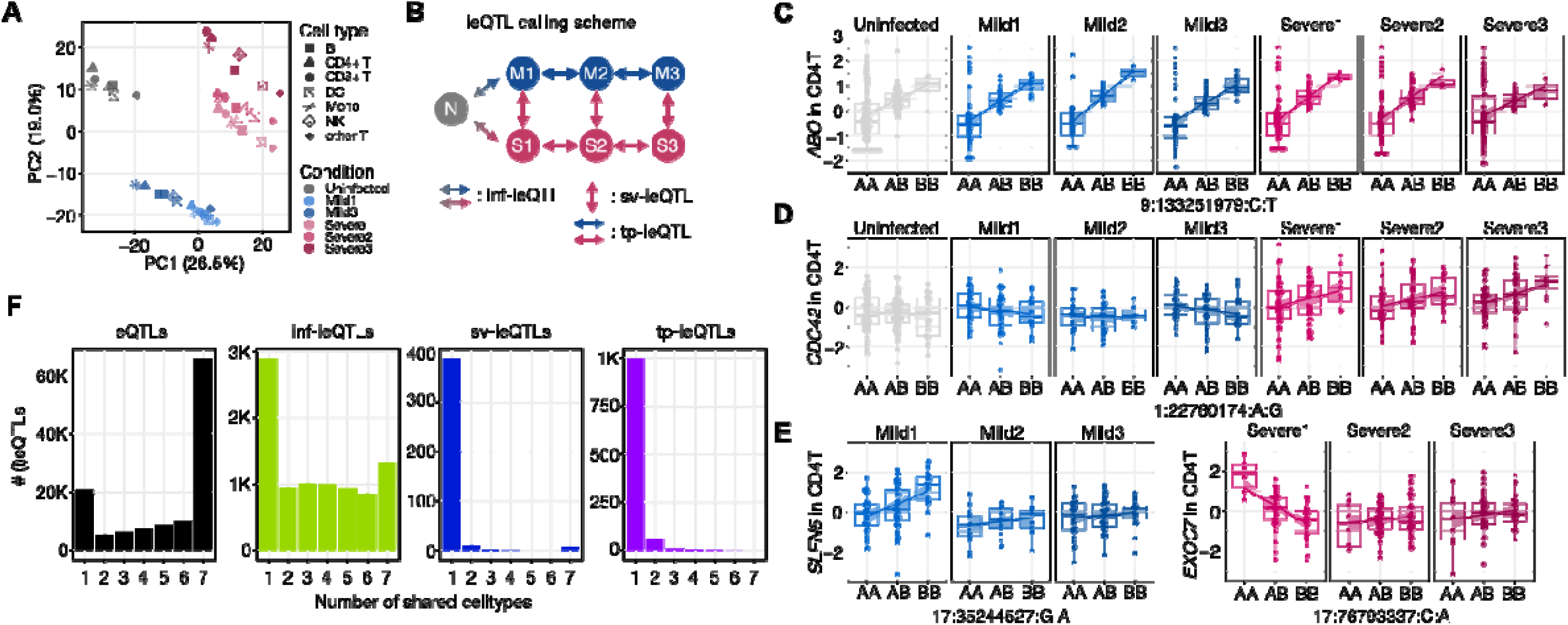
COVID-eQTLs. (A) PCA of eQTLs by disease condition and cell types. (B) ieQTL calling scheme. inf, infection; sv, severity; tp, timepoint. (C) An example eQTL that regulates *ABO* expression. (D) An example sv-ieQTL that regulates *CDC42* expression only in the Severe condition. (E) Example tp-ieQTLs that regulate *SLFN5* and *EXOC7* expression at different timepoints. (F) Number of eQTLs and ieQTLs across cell types.

To investigate eQTL sharing patterns across the cell types, we defined the eQTL with the largest effect size across cell types as the reference and counted the number of cell types in which eQTLs were shared by sign. While a majority of eQTLs recurrently appeared across all major cell types (47,278/89,996 = 52.5%; Fig. 2F, Supplementary Table 5), sv- and tp-ieQTLs showed less overlap, being mostly comprised of unique ieQTLs found in single cell types (number of unique vs. all ieQTLs for sv-ieQTLs: 385/404 = 95.3% and tp-ieQTLs: 904/972 = 93.0%; Fig. 2F, Supplementary Table 6). This finding suggests that ieQTL may function in a more cell-type-specific manner compared to eQTLs, which provide more shared regulatory activities across multiple cell types. Consistent with this, we found ieQTLs also showed lower sharing when quantified from mash posterior estimates rather than by significance, a measure that does not depend on threshold crossing: effects were concordant in direction across all seven cell types for 27.7% of ieQTLs versus 33.5–45.3% of eQTL, and shared by magnitude across all seven for 2.9% versus 4.4–8.6%.

### COVID-eQTLs regulate genes of high constraint and strong expression

Next, we analyzed various genetic features of our COVID-eQTL calls. Plotting the distance of COVID-eQTLs from transcription start sites (TSS) revealed eQTLs to be more centered around TSS compared to ieQTLs, suggesting that the expression regulation posed by ieQTLs is less promoter-centered and therefore more likely to be context-specific (Fig. 3A). When considering COVID-eQTLs according to whether they are common among examined cell types or distinct to a single cell type, we found cell-type-specific COVID-eQTLs to have larger effect sizes than shared COVID-eQTLs with different strengths (eQTLs: two-tailed *t*-test *P* = 9.0 x 10^-148^; ieQTLs: two-tailed *t*-test *P* = 0.00042; Combined *P* = 2.6 x 10^-102^; Fig. 3B and Supplementary Figure 3 A-D). Similarly, when eQTLs were compared by their disease status dependencies, ieQTLs showed smaller effect sizes than eQTLs (two-tailed *t*-test *P* = 1.2 x 10^-134^) (Supplementary Figure 3E). This result implies that regulatory variants which influence cell-type specificity or disease severity through changes in gene expression have a more pronounced influence compared to those found in shared cell-type or disease state. Enrichments of COVID-eQTLs in epigenetic and genomic features across all major cell types further revealed that they are preferentially found in or near TSS, enhancers, 5’UTRs, and exons (Fig. 3C and Supplementary Figure 5). Enrichment analysis revealed that eGenes are enriched in COVID-19 and other immune-related pathways, demonstrating that our calling pipeline has identified a collection of biologically plausible genes (Fig. 3D).

**Figure 3.**
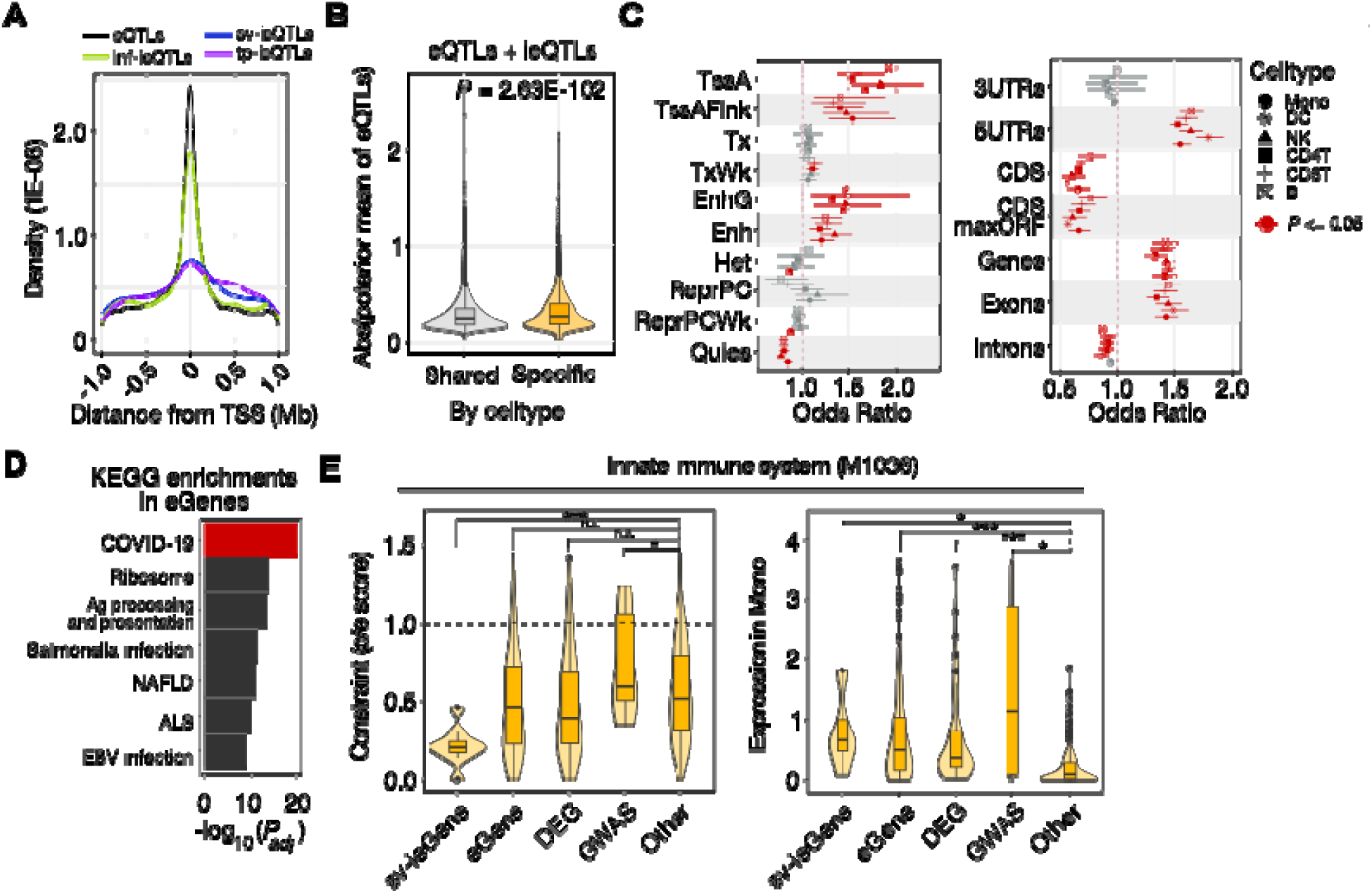
Genetic features of COVID-eQTLs. (A) Distribution of COVID-eQTL positions relative to TSS. (B) Difference in mean β values (*i.e*., effect sizes) between shared and specific eQTLs according to cell types. (C) Enrichment of eQTLs by epigenetic and genomic features. Red dots denote significant enrichment (*P* < 0.05). Error bars represent standard error. TssA, active TSS; TssAFlnk, flanking active TSS; Tx, strong transcription; TxWk, weak transcription; EnhG, genetic enhancer; Enh, enhancer; Het, heterochromatin; ReprPC, repressed polycomb; ReprPCWk, weakly repressed polycomb; Quies, quiescent; CDS maxORF, CDS with maximum ORF. (D) KEGG pathways enriched among eGenes. Ag, antigen; NAFLD, non-alcoholic fatty liver disease; ALS, amyotrophic lateral sclerosis; EBV, Epstein-Barr virus. (E) Distribution of gene constraint scores (LoF o/e score) and expression levels in monocytes (left) and CD8+ T cells (right) by gene categories.

Next, we sought to compare the characteristics of genes that behave as ieGenes, eGenes, differentially expressed genes (DEGs), GWAS-associated genes, or others. We selected two pathways from the Reactome Pathway database[55] that showed significant enrichment for ieGene in monocytes in relation to disease severity based on ToppGene[56] analysis, innate immune system (*P_adj_* = 1.9 x 10^-2^) and adaptive immune system (*P_adj_* = 1.9 x 10^-2^) pathways. Then, we compared the gene constraint scores (o/e constraint score from the gnomAD) and expression levels in monocytes and CD8+ T cells of the genes within each pathway (Fig. 3E). Interestingly, genes whose expression was influenced by ieQTLs (*i.e*., ieGenes) exhibited the highest constraint scores and expression, followed by eGenes, DEGs, and GWAS associated genes. This result was replicated in different cell types, but not in irrelevant pathways (Supplementary Figures 6 and 7). This finding suggests that ieQTLs comprise a preferred method of expression regulation for critical and highly expressed genes; this regulatory effect is context-dependent, and hence avoids ubiquitous large fluctuations in expression.

### Colocalization of COVID-eQTLs with COVID-19 GWAS loci

To test validity of our COVID-eQTL set, we compared our set with previously reported bulk-eQTLs identified from peripheral blood samples (*i.e.*, GTEx blood data and Japan COVID-19 data [23,57]). We observed strong replication of eQTL effect size (Fig. 4A and 4B) for eQTLs, including against a new large-scale single-cell PBMC eQTL dataset (1M-scBloodNL; Fig. 4A), indicating that the majority of variant regulatory effects are shared with other datasets across different genetic ancestries. One of the key advantages of eQTL analysis is colocalization: the eQTLs colocalized with GWAS variants can confer the molecular mechanism underlying the genetic association, through altering expression of nearby genes [58]. Therefore, we sought to determine how many of the loci identified from COVID-19 GWAS were explained by our COVID-eQTL set [6,59]. Overall, we observed a high probability of eQTL colocalization with COVID-19 GWAS signal (PP.H4 ≥ 0.8) in 16 genes, with the greatest number of colocalizations occurring for monocyte eQTLs (Fig. 4C). This level of colocalization aligns with findings from previous COVID-19 GWAS-eQTL colocalization studies in individuals of European ancestries [19–21,60]. Considering that there is only a marginal difference between cell types in terms of the number of COVID-eQTLs called (Supplementary Figure 4), the enrichment of monocyte eQTLs in GWAS variants associated with COVID-19 severity is notable, providing evidence for the importance of the innate immune response in disease severity [50,51,61]. Colocalized eGenes mostly belonged to the *HLA* locus or were previously colocalized genes (Fig. 4D) [17,20,21]. Examples in the *ABO* and *TOMM7* loci are shown in Fig. 4E. This result implies involvement of cell type-specific determination of COVID-19 severity through comparison with GWAS loci. We further conducted a colocalization analysis using GWAS results from immune-related diseases, which revealed eQTLs in B cells and monocytes to harbor the highest numbers of overlapping loci (Supplementary Figure 8).

**Figure 4.**
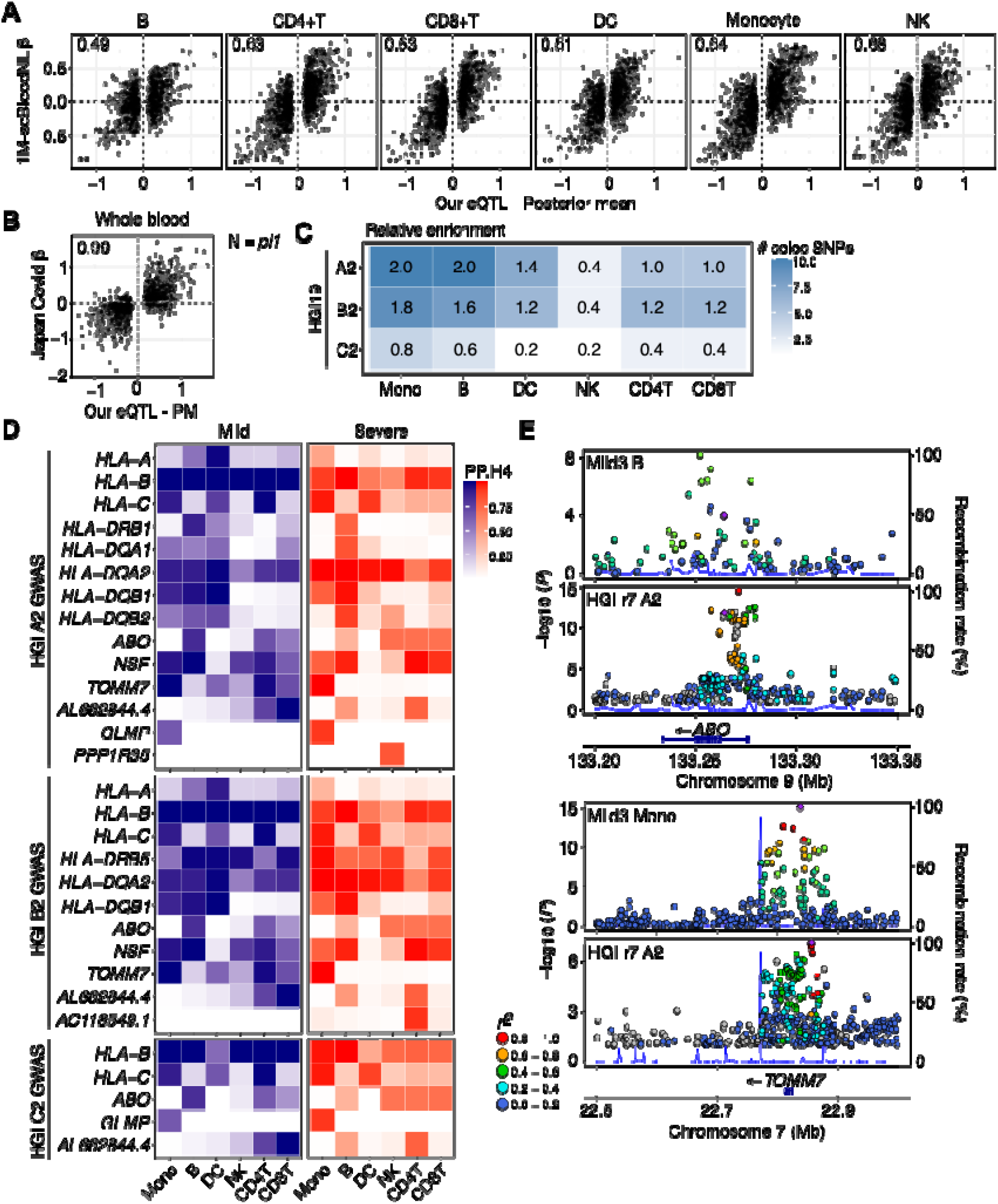
COVID-eQTLs are replicated through exact matches with PBMC eQTLs and colocalization with COVID-19 GWAS. (A) Scatter plot of β values in the COVID-eQTL set versus the 1M-scBloodNL dataset and (B) Japan COV datasets. Number in the upper left corner denotes pi1 value. (C) Heatmap depicting enrichment of colocalization signals for COVID-19 GWAS by cell type. HGI COVID-19 GWAS codes denote as follows: A2, critically ill COVID-19 patients vs. population controls, B2, hospitalized COVID-19 patients vs. non-hospitalized COVID-19 patients, and C2, reported SARS-CoV-2 infected individuals vs. population controls. Values on tiles represent the relative enrichment of colocalized eQTLs for each GWAS phenotype in the given cell types. (D) Heatmap depicting the result of colocalization of eQTLs with A2, B2, and C2. eGenes of colocalized eQTLs are displayed by cell type with the posterior probability H4 (PP.H4) value. (E) Regional plots for two example colocalized loci, showing COVID-eQTL (top) and COVID-19 GWAS (bottom) signals for each locus.

### Genetic regulation of cytokine expression

Levels of circulating cytokines harbor critical utility in the response to viral infection, including in COVID-19 [62–64]. Accordingly we explored blood cytokine levels for protein QTL (pQTL) calling and compared the results with COVID-eQTLs relating to cellular cytokine RNA levels (Supplementary Table 7). Heritability determination revealed similar values for cytokine proteins (mean = 0.04) and transcripts (in the pseudo-bulk level, mean = 0.04) (Fig. 5A and Supplementary Table 8). Notably, cytokines with high heritability were not enriched with COVID-19-related cytokines or DEPs (Wilcoxon rank-sum test *P* = 0.93 and 0.79, respectively; Supplementary Figure 9), nor associated with protein multimerization, post-translational cleavage, or protein half-life (data not shown). Two genes important in the inflammatory response to COVID-19 [65,66], IL6R and MICA, harbored strong pQTLs but non-significant eQTLs (Fig. 5B). Direct comparisons of protein and RNA expression patterns revealed enhanced detection and separation by disease severity for some of the cytokines (Fig. 5C and Supplementary Figure 10). Notably, a region in chromosome 17 encompassing *CCL23* and *CCL18* represents an example of independent regulation by two neighboring pQTLs (Fig. 5D). Conditional analysis revealed that the top SNPs for each protein behaved in a completely independent manner, explaining the observation that these proteins show low correlation of expression despite the encoding genes being close to each other (Supplementary Figure 11).

**Figure 5.**
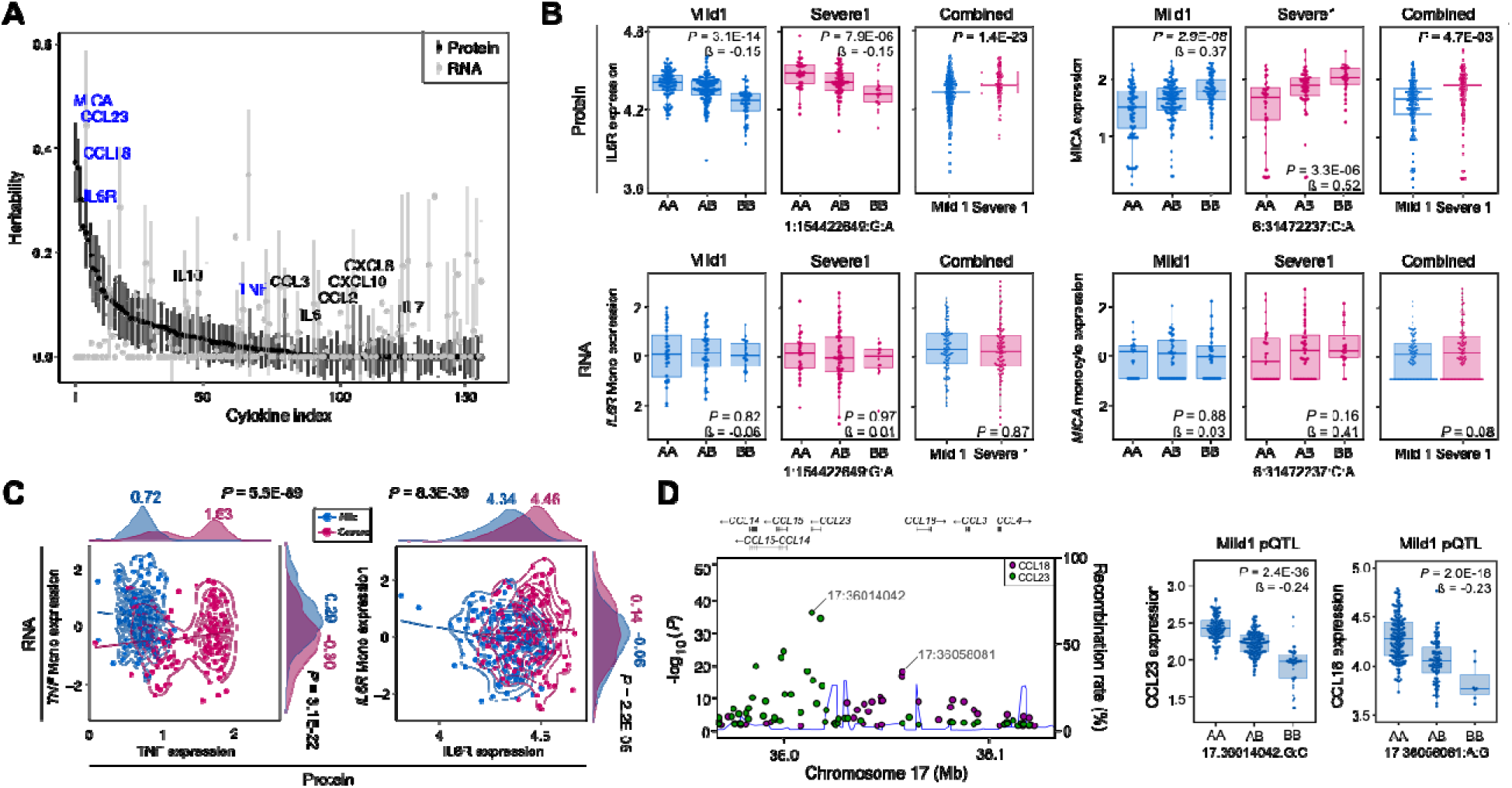
Regulation of blood cytokine production. (A) Distribution of heritability values for cytokine protein level (Protein) or scRNA-seq-based pseudo-bulk cytokine gene expression level (RNA). Cytokines showing different expression between the Mild and Severe groups are marked as DEPs. (B) Cytokines (IL6R and MICA) showing strong pQTLs and weak eQTLs. (C) Scatter plot of TNF and IL6R protein versus RNA expression. (D) A locus in chromosome 17 showing pQTLs with strong but independent effects on CCL23 and CCL18.

### Identification of upstream factors leading to severity-specific eQTLs

It is critical to identify factors that act upstream of many eQTLs as they may represent master regulators that transmit physiological cues in the cell to affect the expression of many downstream genes [25]. However, upstream factors that regulate eQTLs are less studied compared to the downstream effects posed by eQTLs. To examine such factors that influence COVID-19 severity, we devised a model where differences in TF activity (*i.e.*, “regulons”) pose a change in target gene (*i.e*., sv-ieGenes) expression through binding to a regulatory region with differential binding affinity (*i.e.*, ieSNP) (Fig. 6A). Our initial pool was composed of 404 sv-ieQTLs, called from every examined cell type, and 376 TFs, as defined by SCENIC [46], totaling 145,888 regulon-ieQTL pairs. After initial filtering by significant changes in TF-DNA binding ability using FIMO [48], 567 regulon-ieQTL pairs remained (Supplementary Table 9). Then, as a stringent filtering strategy, we applied three criteria to identify regulons that are significantly correlated with (i) differential activity between the Mild and Severe groups, (ii) differential level of downstream gene (ieGene) expression, and (iii) ieQTL effect size, under the assumption that a change of regulon activity in a cell causes a change in ieQTL effect size (Figs. 6B-C). This strategy yielded 18 regulon-ieQTL pairs (Figs. 6B-D), among which we were especially intrigued by the regulation of *IGFBP7* by TF PATZ1 in monocytes. Although its implication in COVID-19 remains elusive, IGFBP7 poses multifaceted functions in inflammatory contexts, such as acute kidney injury and cancers [67,68]. PATZ1 displays significant differential DNA binding ability in relation to the genotype of rs881382 (*i.e.*, a Mild-specific ieSNP in monocytes, Supplementary Figure 12), with the PATZ1 motif containing the reference allele (G) (Fig. 6E). The region flanking rs881382, located in *IGFBP7* intron 1, is an open chromatin region according to ATAC-seq, DNase-seq, and H3K27ac and PATZ1 ChIP-seq from the ENCODE dataset (Fig. 6F and Supplementary Figure 13). Activity of the PATZ1 regulon was higher in the mild group compared to the severe group (two-tailed *t*-test *P* = 3.36 x 10^-21^) (Fig. 6G). Expression of *IGFBP7* showed a negative correlation with PATZ1 as a transcriptional repressor (Pearson’s *R* = -0.39) (Fig. 6H) [69]. And the effect size of the *IGFBP7* eQTL was positively correlated with PATZ1 activity (Pearson’s *R* = 0.79) (Fig. 6I), causing eQTL effect size to be higher in the Mild group (*i.e.*, establishment of a Mild-specific ieQTL) (Fig. 6J). Based on this finding, we propose a model in which differential TF activities in disease status lead to changes in downstream gene expression and thereby constitute a sophisticated gene expression regulatory mechanism (Fig. 6K).

**Figure 6.**
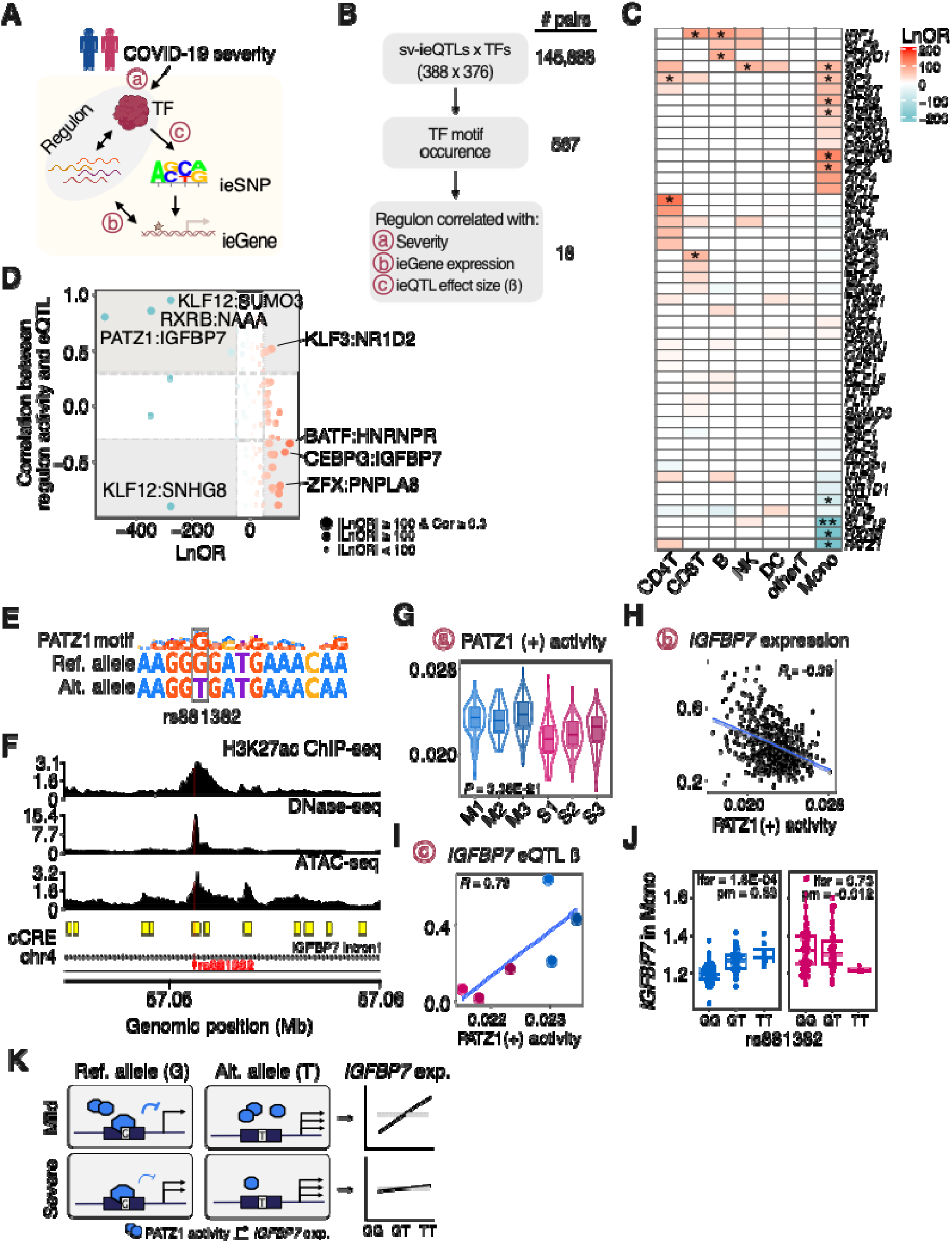
Identification of upstream factors leading to severity-specific eQTLs. (A) Schematic representation of the expected upstream mechanism of ieQTLs. (B) Flow chart displaying a strategy to select strong regulon-ieQTL pairs. (C) Heatmap showing log odds ratio for the candidate regulon-ieQTL pairs. A positive log odds ratio indicates regulons with greater activity in the Severe group, while a negative ratio reflects higher activity in the Mild group. (D) Association of correlation coefficients of regulon activity with ieQTL effect size with log odds ratios (LnOR) for condition-specific regulon activity. Eighteen pairs were highly skewed toward a specific severity group and showed high correlation of regulon activity and eQTL effect size, indicated by grey areas. (E) Binding motifs of PATZ1 along with the observed sequences flanking rs881382. (F) Epigenetic tracks of the region flanking rs881382. (G) PATZ1 regulon activity in monocytes across conditions. (H) Correlation of regulon activity and IGFBP7 expression in monocytes. (I) Correlation of regulon activity and the effect size of the IGFBP7 eQTL. (J) Severity-specific regulation of IGFBP7 expression by the ieQTL under mild conditions. (K) Proposed model for the mild-disease-specific IGFBP7 ieQTL.

## Discussion

Genetic susceptibility of host response to virus infection carries profound implications in both clinical and social systems, and the recent COVID-19 pandemic provided an unprecedented opportunity to systematically address such susceptibility. Indeed, many studies investigated differential genetic and transcriptional profiles in patients with varying disease severity [6,7,19,20,50,59,70]. However, these GWAS variants explain only a limited portion of phenotypic variation, and the rare pathogenic variants are found in only a relatively small portion of patients.

In this study, we used genome, single-cell transcriptome, and blood cytokine data generated from Korean patients infected with COVID-19 to systemically delineate genetic differences between patients with mild or severe responses by calling COVID-eQTLs, or eQTLs and ieQTLs that drive differential gene expression in relation to disease severity and progression. This approach led us to identify 89,996 eQTLs and 1,242 sv- and tp-ieQTLs across seven main cell types of peripheral blood mononuclear cells (PBMCs).

Our findings emphasize the importance and utility of ieQTLs in regulating genes in the context of viral infection. The ieQTL strategy allows consideration of the inherent heterogeneity in cell states among different samples of the same clinical status and hence discovery of variants that regulate gene expressions in a cell-state-dependent manner. This approach was previously utilized in lung and liver diseases to reveal disease-specific regulatory effects and identify pathways associated with cell state [12,25]. Here, we observed genes regulated by ieQTLs to feature both strong constraint and higher expression than genes in other classes, such as those associated with GWAS loci, DEGs, or non-implicated genes (Fig. 3E). This observation suggests that genes under control of eQTLs and ieQTLs may perform more essential functions than GWAS genes in the context of COVID-19 response.

Intriguingly, a number of features that signify disease severity were most prominent in monocytes. Monocytes harbored the largest numbers of eQTLs and ieQTLs (Supplementary Figure 4), eQTLs detected from monocytes demonstrated the highest colocalization with COVID-19-related traits (Fig. 4C), and monocytes exhibited the greatest enrichment of TFs and sv-ieQTL pairs (Fig. 6C). These findings add further evidence of the pivotal role of innate immune cells in cytokine production and the determination of COVID-19 severity [10,51,53,70].

Current understanding of the genetic regulation of cytokine levels is limited. Previous studies have reported high pathogen-dependent variability when cells are stimulated, including by virus [71–73]. Our quantitative comparison of genetic influence on cytokine protein expression in blood and gene expression in PBMCs revealed genetic factors to have similar ability to predict RNA and cytokine levels but the degree of dependency on either RNA- or protein-level differs by each cytokine.

One advantage of eQTL approach is that it can instantly identify pathways whose activity may be dependent on individuals’ genotypes. Another is that it enables the elucidation of immediate upstream factors, such as TFs, that directly regulates many eQTLs. IGFBP7 is a versatile secreted protein with roles in acute kidney injury, cancers, and vascular diseases, which it enacts through regulating fibrosis, inflammation, senescence, and oxidative stress [68,74]. It is expressed in many cell types of PBMCs and implicated as an upstream regulator of genes important in inflammation [75,76]. Expression of *IGFBP7* is considered dynamic, with controversial reports supporting different patterns of expression depending on the disease model [76–78]. Through probing a regulatory network involving the ieQTL and upstream regulators of *IGFBP7* expression, we suggest that PATZ1, a transcription factor whose activity differs by disease severity, together with the allele status of rs881382, located in the intron 1 of *IGFBP7*, contributes to the cell-state-specific regulation of *IGFBP7*. We envision that this approach will be used to accurately understand the behavior of many interesting ieQTLs in the future.

While our approach mainly considered the systematic combination of genome and transcriptomic data, investigating the epigenetic regulation of these genes in COVID-19 patients may provide additional mechanistic insights into differential disease severity. For example, probing of open chromatin regions in hospitalized COVID-19 patients revealed an enrichment of CEBP TFs in monocytes, which aligns with our findings [79]. In addition, more fine stratification of cell states (*e.g.*, gene modules) using a larger cohort may provide further insights into the regulation of genes whose expression relates to disease severity.

In summary, we combined extensive multimodal data from COVID-19 patients and identified differential regulation of gene expression in relation to disease severity across cell types. Our COVID-eQTL analysis pipeline will serve as a novel framework for elucidating the biological determination of disease severity and as a tool for discovering predictive biomarkers, not only for SARS-CoV-2, but also for other future viral pandemics.

## Conclusions

We combined genomic, single-cell transcriptomic and cytokine data collected over time from Korean COVID-19 patients to map how genetic variation shapes immune gene expression across disease severity and progression. Mapping eQTLs with the disease state taken into account identified regulatory effects that depend on cell type and context. It also linked these effects to known COVID-19 risk loci and to the transcription factors that act upstream of them. The framework reveals statistical association to identify mechanisms of host genetic susceptibility. It can be applied to other infectious diseases and to future pandemics.

## List of abbreviations

COVID-19: Coronavirus disease 2019
eGene: Gene regulated by an eQTL
eQTL: Expression quantitative trait locus
eSNP: SNP associated with gene expression (eQTL variant)
GWAS: Genome-wide association study
HGI: COVID-19 Host Genetics Initiative
HLA: Human leukocyte antigen
ieGene: Gene regulated by an ieQTL
ieQTL: Interaction expression quantitative trait locus
ieSNP: SNP of an ieQTL
inf-ieQTL: Infection-interacting ieQTL
ISG: Interferon-stimulated gene
KOVA: Korean Variant Archive
LFSR: Local false sign rate
MAF: Minor allele frequency
PBMC: Peripheral blood mononuclear cell
PCA: Principal component analysis
pQTL: Protein quantitative trait locus
SARS-CoV-2: Severe acute respiratory syndrome coronavirus 2
scRNA-seq: Single-cell RNA sequencing
sv-ieQTL: Severity-interacting ieQTL
tp-ieQTL: Timepoint-interacting ieQTL

## Acknowledgements

We appreciate participation of the COVID-19 patients and clinicians involved in the study. We thank Jungmin Choi, Seunghee Hong, Joon Yong An, Yongjin Yoo, and Sung Eun Hong for their critical reading of the manuscript.

## Data availability

All clinical data and biospecimens from this project are available from the National Biobank of Korea (NBK) under prior approval (https://nih.go.kr/biobank). Also, omics dataset from this project are deposited at the Clinical and Omics data Archive (CODA; https://coda.nih.go.kr) via accession numbers CODA_D21001 and CODA_D23017.

## Code availability

Code for the analysis is available at https://github.com/snu-mchoi-lab/COVID-QTL. Most analyses are performed using R. The open-source tools used for data analysis are referred to in Methods or Supplementary Methods.

## Author information

### Contributions

W.-Y.P., M.C., S.Z. conceived the study. J.L., E.Y.J., M.C. designed the experiments and analyses. S.Z., M.C. acquired funding and provided supervision. J.L., E.Y.J. performed scRNA-seq data analysis, genome data processing, eQTL calling, and downstream analyses. L.Y., G.D., S.Z. called interaction-eQTLs and performed downstream analyses. J.L., E.Y.J., L.Y., S.Z., M.C. wrote the paper. H.-Y.J., S.C.K., W.- Y.P., H.-Y.P. contributed to data generation, preprocessing, and administrative tasks.

### Corresponding authors

These authors jointly supervised this work: Siming Zhao, Murim Choi

## Ethics declarations

The authors declare no conflicts of interest that pertain to this work.

## References

1. Kwok AJ, Mentzer A, Knight JC. Host genetics and infectious disease: new tools, insights and translational opportunities. Nat Rev Genet. 2021;22:137–53.

2. O’Driscoll M, Ribeiro Dos Santos G, Wang L, Cummings DAT, Azman AS, Paireau J, et al. Age-specific mortality and immunity patterns of SARS-CoV-2. Nature. 2021;590:140–5.

3. Takahashi T, Ellingson MK, Wong P, Israelow B, Lucas C, Klein J, et al. Sex differences in immune responses that underlie COVID-19 disease outcomes. Nature. 2020;588:315–20.

4. Zhou F, Yu T, Du R, Fan G, Liu Y, Liu Z, et al. Clinical course and risk factors for mortality of adult inpatients with COVID-19 in Wuhan, China: a retrospective cohort study. Lancet. 2020;395:1054–62.

5. Docherty AB, Harrison EM, Green CA, Hardwick HE, Pius R, Norman L, et al. Features of 20 133 UK patients in hospital with covid-19 using the ISARIC WHO Clinical Characterisation Protocol: prospective observational cohort study. BMJ. 2020;369:m1985.

6. Initiative C-HG. A second update on mapping the human genetic architecture of COVID-19. Nature. 2023;621:E7–E26.

7. Zhang Q, Bastard P, Effort CHG, Cobat A, Casanova JL. Human genetic and immunological determinants of critical COVID-19 pneumonia. Nature. 2022;603:587–98.

8. Zhang Q, Bastard P, Liu Z, Le Pen J, Moncada-Velez M, Chen J, et al. Inborn errors of type I IFN immunity in patients with life-threatening COVID-19. Science. 2020;370.

9. Li Y, Ke Y, Xia X, Wang Y, Cheng F, Liu X, et al. Genome-wide association study of COVID-19 severity among the Chinese population. Cell Discov. 2021;7:76.

10. Edahiro R, Shirai Y, Takeshima Y, Sakakibara S, Yamaguchi Y, Murakami T, et al. Single-cell analyses and host genetics highlight the role of innate immune cells in COVID-19 severity. Nat Genet. 2023;55:753–67.

11. Vosa U, Claringbould A, Westra HJ, Bonder MJ, Deelen P, Zeng B, et al. Large-scale cis- and trans-eQTL analyses identify thousands of genetic loci and polygenic scores that regulate blood gene expression. Nat Genet. 2021;53:1300–10.

12. Natri HM, Del Azodi CB, Peter L, Taylor CJ, Chugh S, Kendle R, et al. Cell-type-specific and disease-associated expression quantitative trait loci in the human lung. Nat Genet. 2024;56:595–604.

13. Yoo T, Joo SK, Kim HJ, Kim HY, Sim H, Lee J, et al. Disease-specific eQTL screening reveals an anti-fibrotic effect of AGXT2 in non-alcoholic fatty liver disease. J Hepatol. 2021;75:514–23.

14. Hong SE, Choi M, Lee J. Role of expression quantitative trait loci (eQTL) in understanding genetic mechanisms underlying common complex diseases. Mol Cells. 2025;48:100256.

15. Consortium GT. The Genotype-Tissue Expression (GTEx) project. Nat Genet. 2013;45:580–5.

16. Yazar S, Alquicira-Hernandez J, Wing K, Senabouth A, Gordon MG, Andersen S, et al. Single-cell eQTL mapping identifies cell type-specific genetic control of autoimmune disease. Science. 2022;376:eabf3041.

17. D’Antonio M, Nguyen JP, Arthur TD, Matsui H, Initiative C-HG, D’Antonio-Chronowska A, et al. SARS-CoV-2 susceptibility and COVID-19 disease severity are associated with genetic variants affecting gene expression in a variety of tissues. Cell Rep. 2022;39:110968.

18. Baranova A, Cao H, Zhang F. Unraveling Risk Genes of COVID-19 by Multi-Omics Integrative Analyses. Front Med (Lausanne). 2021;8:738687.

19. D’Antonio M, Nguyen JP, Arthur TD, Matsui H, Initiative C-HG, D’Antonio-Chronowska A, et al. SARS-CoV-2 susceptibility and COVID-19 disease severity are associated with genetic variants affecting gene expression in a variety of tissues. Cell Rep. 2021;37:110020.

20. Schmiedel BJ, Rocha J, Gonzalez-Colin C, Bhattacharyya S, Madrigal A, Ottensmeier CH, et al. COVID-19 genetic risk variants are associated with expression of multiple genes in diverse immune cell types. Nat Commun. 2021;12:6760.

21. Aquino Y, Bisiaux A, Li Z, O’Neill M, Mendoza-Revilla J, Merkling SH, et al. Dissecting human population variation in single-cell responses to SARS-CoV-2. Nature. 2023;621:120–8.

22. Randolph HE, Aguirre-Gamboa R, Brunet-Ratnasingham E, Nakanishi T, Mu Z, Locher V, et al. Widespread gene-environment interactions shape the immune response to SARS-CoV-2 infection in hospitalized COVID-19 patients. Nat Commun. 2026;17.

23. Wang QS, Edahiro R, Namkoong H, Hasegawa T, Shirai Y, Sonehara K, et al. The whole blood transcriptional regulation landscape in 465 COVID-19 infected samples from Japan COVID-19 Task Force. Nat Commun. 2022;13:4830.

24. Mostafavi H, Spence JP, Naqvi S, Pritchard JK. Systematic differences in discovery of genetic effects on gene expression and complex traits. Nat Genet. 2023;55:1866–75.

25. Hong SE, Mun SJ, Lee YJ, Yoo T, Suh KS, Kang KW, et al. Single-cell eQTL analysis identifies genetic variation underlying metabolic dysfunction-associated steatohepatitis. Nat Genet. 2025;57:1638–48.

26. Jo HY, Kim SC, Ahn DH, Lee S, Chang SH, Jung SY, et al. Establishment of the large-scale longitudinal multi-omics dataset in COVID-19 patients: data profile and biospecimen. BMB Rep. 2022;55:465–71.

27. COVID-19 Treatment Guidelines Panel. Coronavirus Disease 2019 (COVID-19) Treatment Guidelines. National Institutes of Health. Available at https://www.covid19treatmentguidelines.nih.gov/. Accessed [6/26/2024].

28. Wolock SL, Lopez R, Klein AM. Scrublet: Computational Identification of Cell Doublets in Single-Cell Transcriptomic Data. Cell Syst. 2019;8:281–91 e9.

29. Wolf FA, Angerer P, Theis FJ. SCANPY: large-scale single-cell gene expression data analysis. Genome Biol. 2018;19:15.

30. Hao Y, Stuart T, Kowalski MH, Choudhary S, Hoffman P, Hartman A, et al. Dictionary learning for integrative, multimodal and scalable single-cell analysis. Nat Biotechnol. 2024;42:293–304.

31. Korsunsky I, Millard N, Fan J, Slowikowski K, Zhang F, Wei K, et al. Fast, sensitive and accurate integration of single-cell data with Harmony. Nat Methods. 2019;16:1289–96.

32. Hao Y, Hao S, Andersen-Nissen E, Mauck WM, 3rd, Zheng S, Butler A, et al. Integrated analysis of multimodal single-cell data. Cell. 2021;184:3573–87 e29.

33. DePristo MA, Banks E, Poplin R, Garimella KV, Maguire JR, Hartl C, et al. A framework for variation discovery and genotyping using next-generation DNA sequencing data. Nat Genet. 2011;43:491–8.

34. Li H. Toward better understanding of artifacts in variant calling from high-coverage samples. Bioinformatics. 2014;30:2843–51.

35. Lee S, Seo J, Park J, Nam JY, Choi A, Ignatius JS, et al. Korean Variant Archive (KOVA): a reference database of genetic variations in the Korean population. Sci Rep. 2017;7:4287.

36. Zhou HJ, Li L, Li Y, Li W, Li JJ. PCA outperforms popular hidden variable inference methods for molecular QTL mapping. Genome Biol. 2022;23:210.

37. Taylor-Weiner A, Aguet F, Haradhvala NJ, Gosai S, Anand S, Kim J, et al. Scaling computational genomics to millions of individuals with GPUs. Genome Biol. 2019;20:228.

38. Urbut SM, Wang G, Carbonetto P, Stephens M. Flexible statistical methods for estimating and testing effects in genomic studies with multiple conditions. Nat Genet. 2019;51:187–95.

39. Giambartolomei C, Vukcevic D, Schadt EE, Franke L, Hingorani AD, Wallace C, et al. Bayesian test for colocalisation between pairs of genetic association studies using summary statistics. PLoS Genet. 2014;10:e1004383.

40. Wang G, Sarkar A, Carbonetto P, Stephens M. A simple new approach to variable selection in regression, with application to genetic fine mapping. Journal of the Royal Statistical Society Series B: Statistical Methodology. 2020;82:1273–300.

41. Yu G, Wang LG, Han Y, He QY. clusterProfiler: an R package for comparing biological themes among gene clusters. OMICS. 2012;16:284–7.

42. Delaneau O, Ongen H, Brown AA, Fort A, Panousis NI, Dermitzakis ET. A complete tool set for molecular QTL discovery and analysis. Nat Commun. 2017;8:15452.

43. Roadmap Epigenomics C, Kundaje A, Meuleman W, Ernst J, Bilenky M, Yen A, et al. Integrative analysis of 111 reference human epigenomes. Nature. 2015;518:317–30.

44. Frankish A, Diekhans M, Ferreira AM, Johnson R, Jungreis I, Loveland J, et al. GENCODE reference annotation for the human and mouse genomes. Nucleic Acids Res. 2019;47:D766–D73.

45. Yang J, Lee SH, Goddard ME, Visscher PM. GCTA: a tool for genome-wide complex trait analysis. Am J Hum Genet. 2011;88:76–82.

46. Aibar S, Gonzalez-Blas CB, Moerman T, Huynh-Thu VA, Imrichova H, Hulselmans G, et al. SCENIC: single-cell regulatory network inference and clustering. Nat Methods. 2017;14:1083–6.

47. Hie B, Cho H, DeMeo B, Bryson B, Berger B. Geometric Sketching Compactly Summarizes the Single-Cell Transcriptomic Landscape. Cell Syst. 2019;8:483–93 e7.

48. Grant CE, Bailey TL, Noble WS. FIMO: scanning for occurrences of a given motif. Bioinformatics. 2011;27:1017–8.

49. Sloan CA, Chan ET, Davidson JM, Malladi VS, Strattan JS, Hitz BC, et al. ENCODE data at the ENCODE portal. Nucleic Acids Res. 2016;44:D726–32.

50. Stephenson E, Reynolds G, Botting RA, Calero-Nieto FJ, Morgan MD, Tuong ZK, et al. Single-cell multi-omics analysis of the immune response in COVID-19. Nat Med. 2021;27:904–16.

51. Ren X, Wen W, Fan X, Hou W, Su B, Cai P, et al. COVID-19 immune features revealed by a large-scale single-cell transcriptome atlas. Cell. 2021;184:1895–913 e19.

52. Wilk AJ, Rustagi A, Zhao NQ, Roque J, Martinez-Colon GJ, McKechnie JL, et al. A single-cell atlas of the peripheral immune response in patients with severe COVID-19. Nat Med. 2020;26:1070–6.

53. Hadjadj J, Yatim N, Barnabei L, Corneau A, Boussier J, Smith N, et al. Impaired type I interferon activity and inflammatory responses in severe COVID-19 patients. Science. 2020;369:718–24.

54. Combes AJ, Courau T, Kuhn NF, Hu KH, Ray A, Chen WS, et al. Global absence and targeting of protective immune states in severe COVID-19. Nature. 2021;591:124–30.

55. Milacic M, Beavers D, Conley P, Gong C, Gillespie M, Griss J, et al. The Reactome Pathway Knowledgebase 2024. Nucleic Acids Res. 2024;52:D672–D8.

56. Chen J, Bardes EE, Aronow BJ, Jegga AG. ToppGene Suite for gene list enrichment analysis and candidate gene prioritization. Nucleic Acids Res. 2009;37:W305–11.

57. Consortium GT. The GTEx Consortium atlas of genetic regulatory effects across human tissues. Science. 2020;369:1318–30.

58. Nicolae DL, Gamazon E, Zhang W, Duan S, Dolan ME, Cox NJ. Trait-associated SNPs are more likely to be eQTLs: annotation to enhance discovery from GWAS. PLoS Genet. 2010;6:e1000888.

59. Initiative C-HG. Mapping the human genetic architecture of COVID-19. Nature. 2021;600:472–7.

60. Ota M, Nagafuchi Y, Hatano H, Ishigaki K, Terao C, Takeshima Y, et al. Dynamic landscape of immune cell-specific gene regulation in immune-mediated diseases. Cell. 2021;184:3006–21 e17.

61. Martinez FO, Combes TW, Orsenigo F, Gordon S. Monocyte activation in systemic Covid-19 infection: Assay and rationale. EBioMedicine. 2020;59:102964.

62. Jing X, Xu M, Song D, Yue T, Wang Y, Zhang P, et al. Association between inflammatory cytokines and anti-SARS-CoV-2 antibodies in hospitalized patients with COVID-19. Immun Ageing. 2022;19:12.

63. Calvo-Alvarez E, D’Alessandro S, Zanotta N, Basilico N, Parapini S, Signorini L, et al. Multiplex array analysis of circulating cytokines and chemokines in COVID-19 patients during the first wave of the SARS-CoV-2 pandemic in Milan, Italy. BMC Immunol. 2024;25:49.

64. Liu QQ, Cheng A, Wang Y, Li H, Hu L, Zhao X, et al. Cytokines and their relationship with the severity and prognosis of coronavirus disease 2019 (COVID-19): a retrospective cohort study. BMJ Open. 2020;10:e041471.

65. Farzad F, Yaghoubi N, Jabbari-Azad F, Mahmoudi M, Mohammadi M. Prognostic Value of Serum MICA Levels as a Marker of Severity in COVID-19 Patients. Immunol Invest. 2022;51:1856–66.

66. Group RC. Baricitinib in patients admitted to hospital with COVID-19 (RECOVERY): a randomised, controlled, open-label, platform trial and updated meta-analysis. Lancet. 2022;400:359–68.

67. Yu JT, Hu XW, Yang Q, Shan RR, Zhang Y, Dong ZH, et al. Insulin-like growth factor binding protein 7 promotes acute kidney injury by alleviating poly ADP ribose polymerase 1 degradation. Kidney Int. 2022;102:828–44.

68. Zhang L, Lian R, Zhao J, Feng X, Ye R, Pan L, et al. IGFBP7 inhibits cell proliferation by suppressing AKT activity and cell cycle progression in thyroid carcinoma. Cell Biosci. 2019;9:44.

69. Abramova A, Sakaguchi S, Schebesta A, Hassan H, Boucheron N, Valent P, et al. The transcription factor MAZR preferentially acts as a transcriptional repressor in mast cells and plays a minor role in the regulation of effector functions in response to FcepsilonRI stimulation. PLoS One. 2013;8:e77677.

70. Yao C, Bora SA, Parimon T, Zaman T, Friedman OA, Palatinus JA, et al. Cell-type-specific immune dysregulation in severely ill COVID-19 patients. Cell Rep. 2021;34:108943.

71. Brodin P, Jojic V, Gao T, Bhattacharya S, Angel CJ, Furman D, et al. Variation in the human immune system is largely driven by non-heritable influences. Cell. 2015;160:37– 47.

72. Li Y, Oosting M, Smeekens SP, Jaeger M, Aguirre-Gamboa R, Le KTT, et al. A Functional Genomics Approach to Understand Variation in Cytokine Production in Humans. Cell. 2016;167:1099–110 e14.

73. Li Y, Oosting M, Deelen P, Ricano-Ponce I, Smeekens S, Jaeger M, et al. Inter-individual variability and genetic influences on cytokine responses to bacteria and fungi. Nat Med. 2016;22:952–60.

74. Wang X, Li Y, Zhao Z, Meng Y, Bian J, Bao R, et al. IGFBP7 regulates sepsis-induced epithelial-mesenchymal transition through ERK1/2 signaling. Acta Biochim Biophys Sin (Shanghai). 2019;51:799–806.

75. Chen L, Hui L, Li J. The multifaceted role of insulin-like growth factor binding protein 7. Front Cell Dev Biol. 2024;12:1420862.

76. Katoh M, Nomura S, Yamada S, Ito M, Hayashi H, Katagiri M, et al. Vaccine Therapy for Heart Failure Targeting the Inflammatory Cytokine Igfbp7. Circulation. 2024;150:374–89.

77. Mo W, Deng L, Cheng Y, Ge S, Wang J. IGFBP7 regulates cell proliferation and migration through JAK/STAT pathway in gastric cancer and is regulated by DNA and RNA methylation. J Cell Mol Med. 2024;28:e70080.

78. Qiu L, Huang Z. Downregulation of IGFBP7 Alleviates LPS-induced Inflammation and Apoptosis in WI-38 Cells via Enhancing Mitophagy. Cell Biochem Biophys. 2024; doi:10.1007/s12013-024-01567-4.

79. Zhang B, Zhang Z, Koeken V, Kumar S, Aillaud M, Tsay HC, et al. Altered and allele-specific open chromatin landscape reveals epigenetic and genetic regulators of innate immunity in COVID-19. Cell Genom. 2023;3:100232.

