## Supplemental figures for "Integrated analysis of COVID-19 multi-omics data for eQTLs reveals genetic mechanisms underlying disease severity"

**Contents:**

**Supplementary Table 1.** Clinical characteristics of the participants.

**Supplementary Table 2.** Sample sizes for QTL analyses.

**Supplementary Table 3.** List of eQTLs across the six conditions and seven cell types.

**Supplementary Table 4.** List of sv-ieQTLs and tp-ieQTLs.

**Supplementary Table 5.** Numbers and proportions of each cell type cluster.

**Supplementary Table 6.** List of cytokine pQTLs.

**Supplementary Table 7.** Data points from GCTA analysis used in Figure 5A.

**Supplementary Table 8.** List of 567 regulon-ieQTL pairs.

**Supplementary Table 9.** List of sources of other immune diseases for coloc analysis.

**Supplementary Figure 1.** Population PCA of the participants based on variants from WGS.

**Supplementary Figure 2.** ISG expression patterns of each severity category across cell types.

**Supplementary Figure 3.** Effect sizes of eQTLs and ieQTLs classified by the number of shared cell types by sign.

**Supplementary Figure 4.** Counts of eQTLs (left) and ieQTLs across cell types.

**Supplementary Figure 5.** Enrichment of ieQTLs by epigenetic (left) and genomic (right) features.

**Supplementary Figure 6.** Distribution of gene constraint and expression of genes, which are members of innate or adaptive immune system pathways, across cell types.

**Supplementary Figure 7.** Distribution of gene constraint and expression of genes, which are members of cell cycle, muscle contraction, reproduction, and metabolism of lipids pathway, across the cell types.

**Supplementary Figure 8.** Heatmap depicting enrichment of colocalization signals for autoimmune diseases by cell type.

**Supplementary Figure 9.** Volcano plots of differently expressed cytokine proteins from the comparison of M1 vs. S1 and M3 vs. S3.

**Supplementary Figure 10.** Scatter plots of cytokines in relation to COVID-19 pathway (ID: WP4891) with their RNA expression levels.

**Supplementary Figure 11.** Conditional analysis on pQTLs for CCL23 and CCL18.

**Supplementary Figure 12.** Monocyte-specific regulation of *IGFBP7* expression mediating PATZ1 regulon.

**Supplementary Figure 13.** ChIP-seq data for PATZ1 in the IGFBP7 locus in the K562 cell line.

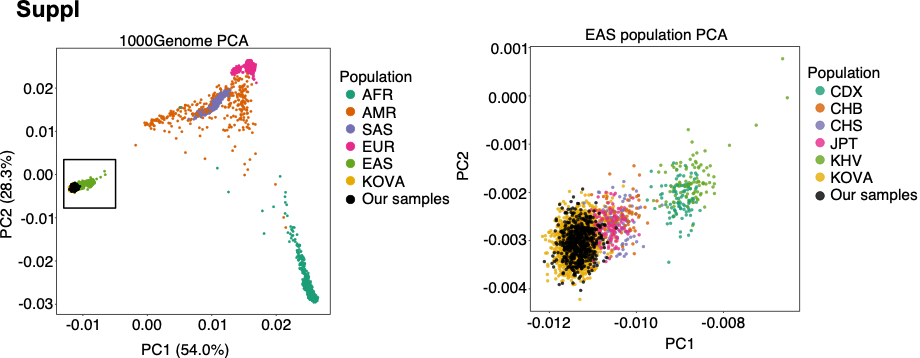

**Supplementary Figure 1.** Population PCA of the participants based on variants from WGS. Population stratification is shown for all ancestries (left) and the East Asian population (right). The East Asian population is highlighted with a black rectangle in the left figure.

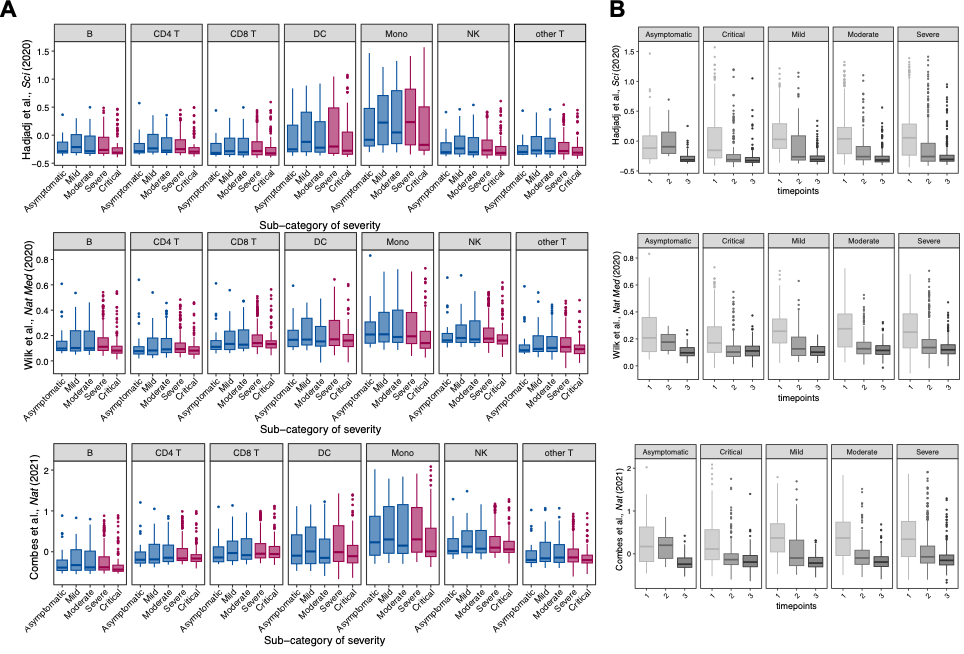

**Supplementary Figure 2.** ISG expression patterns of each severity category across cell types. ISG groups were defined by different studies, as indicated on the left side of each plot. (A) ISG expression in severity category by cell type. (B) ISG expression in time points by severity category.

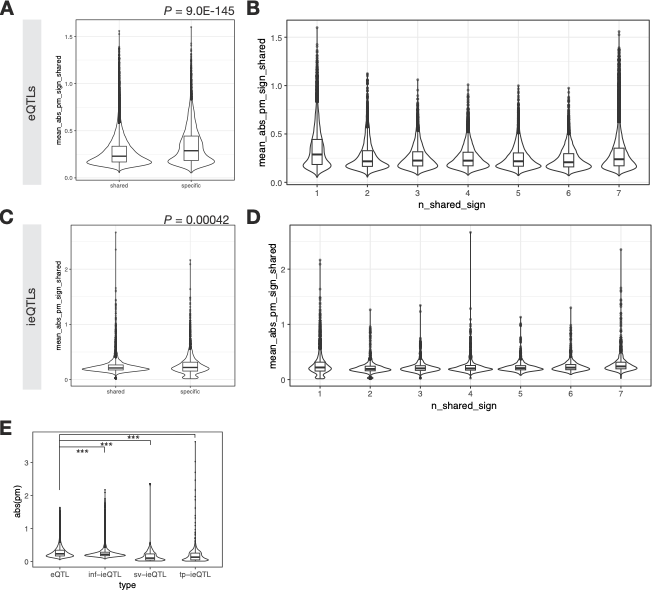

**Supplementary Figure 3.** Effect sizes of eQTLs and ieQTLs classified by the number of shared cell types by sign. (E) shows effect size distribution of eQTLs, inf-ieQTLs, sv-ieQTLs, and tp-ieQTLs (two-sided T test p values between eQTLs vs inf-ieQTL, eQTLS vs sv-ieQTLs, and eQTLs vs tp-ieQTLs are 4.78E-96, 4.60E-13, and 2.13E-40, respectively.

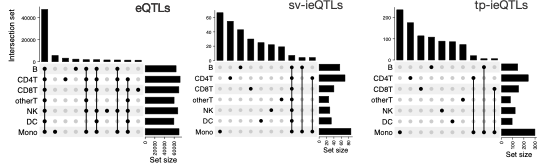

**Supplementary Figure 4.** Counts of eQTLs (left), sv-ieQTLs (middle) and tp-ieQTLs (right) across cell types.

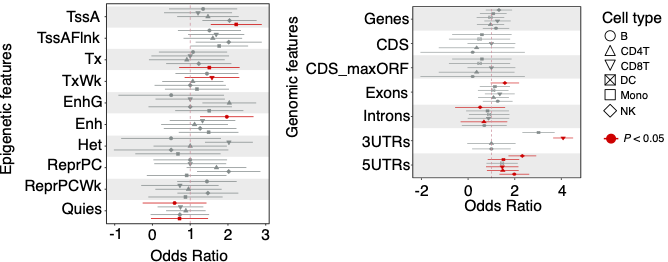

**Supplementary Figure 5.** Enrichment of ieQTLs by epigenetic (left) and genomic (right) features. Error bars represent standard error.

**A**

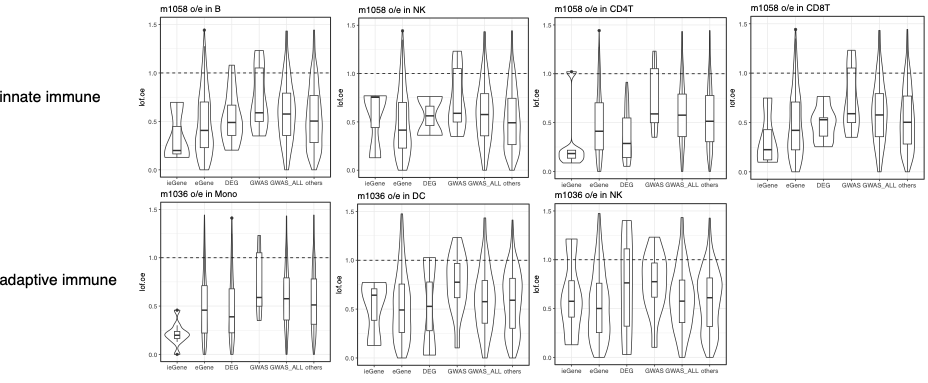

**B**

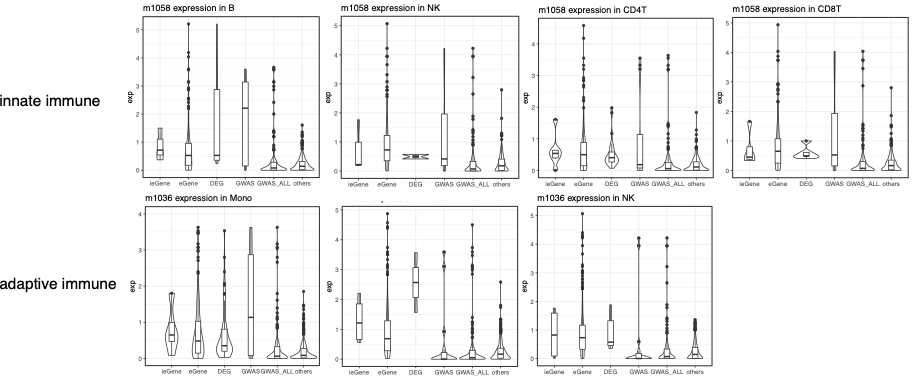

**Supplementary Figure 6.** Distribution of gene constraint and expression of genes, which are members of innate or adaptive immune system pathways, in non-monocytes. (A) Gene constraint shown as o/e scores of genes in the innate immune pathway from NK cells, monocytes, and DCs (upper) and genes in the adaptive immune pathway from CD4 T, CD8 T, NK, and B cells (lower). (B) Expression level of genes in the innate immune pathway from NK cells, monocytes, and DCs (upper) and genes in the adaptive immune pathway from CD4 T, CD8 T, NK, and B cells (lower). Genes in the “GWAS” class denote COVID-19 GWAS-associated genes in the designated pathway. Genes in the “GWAS_ALL” class denote all COVID-19 GWAS-associated genes.

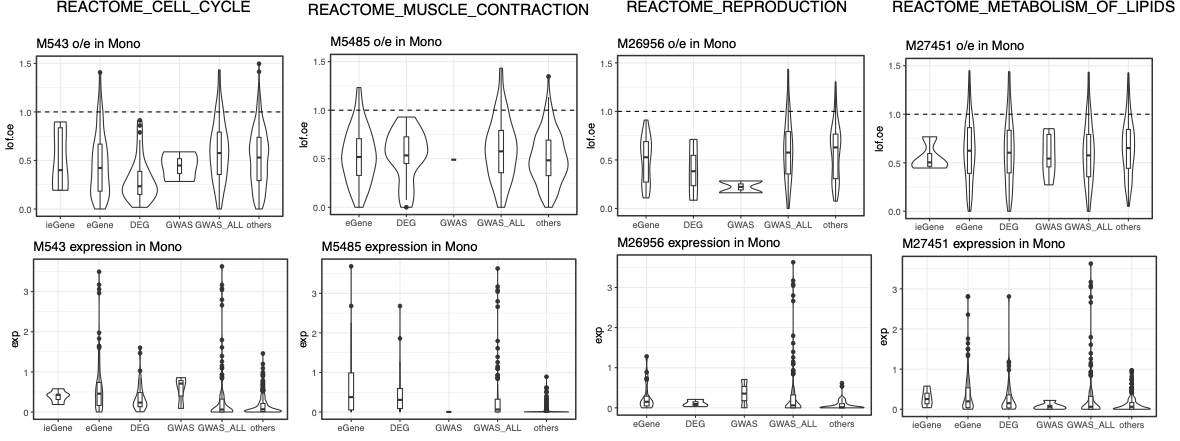

**Supplementary Figure 7.** Distribution of gene constraint and expression of genes, which are members of cell cycle, muscle contraction, reproduction, and metabolism of lipids pathways, in monocytes.

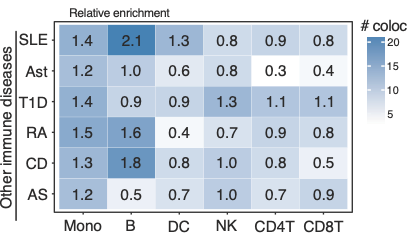

**Supplementary Figure 8.** Heatmap depicting enrichment of colocalization signals for autoimmune diseases by cell type. SLE, systemic lupus erythematosus; Ast, asthma; T1D, type 1 diabetes; RA, rheumatoid arthritis; CD, Crohn’s disease; AS, ankylosing spondylitis.

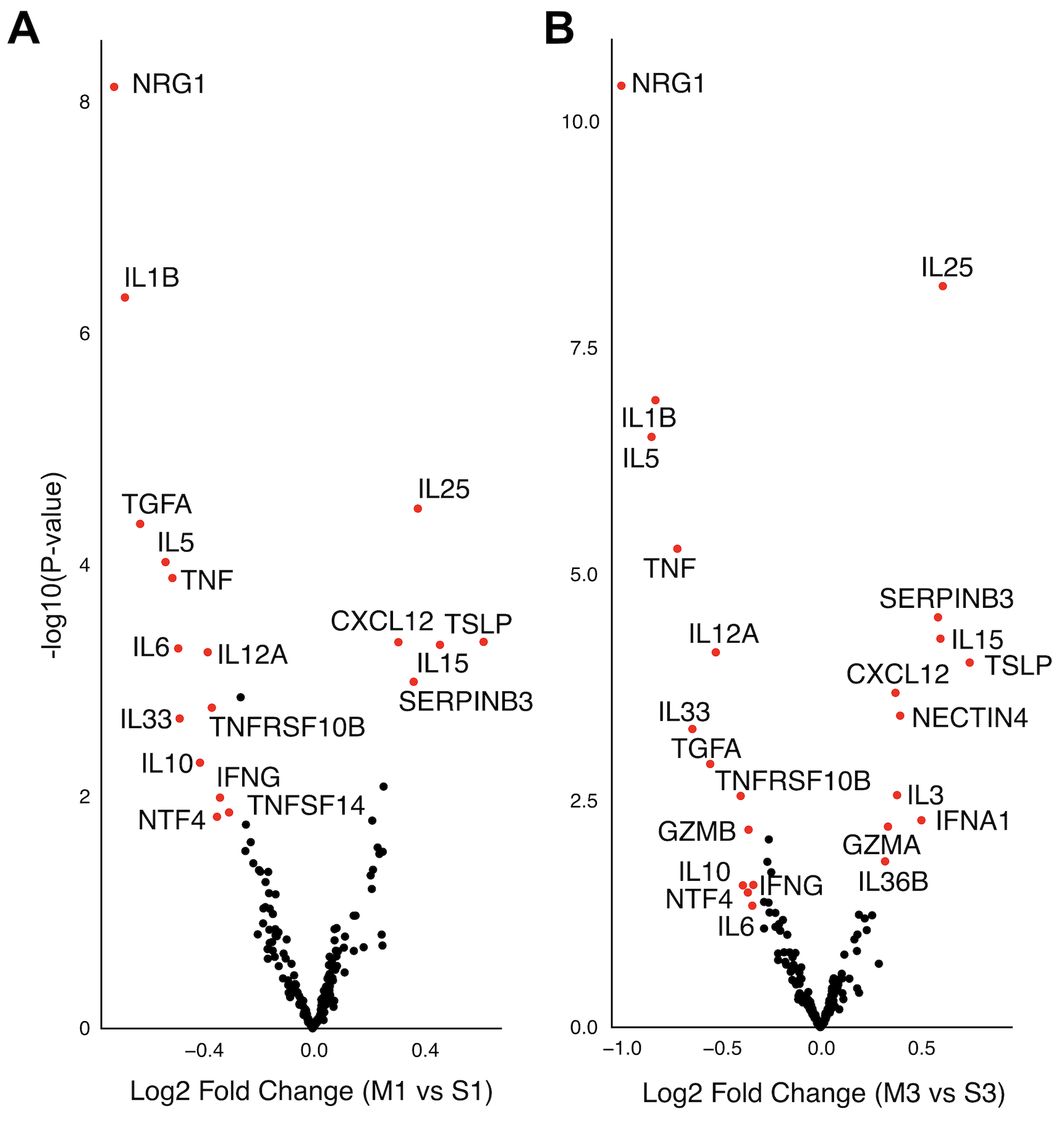

**Supplementary Figure 9.** Volcano plots of differently expressed cytokine proteins from the comparison of M1 vs. S1 and M3 vs. S3.

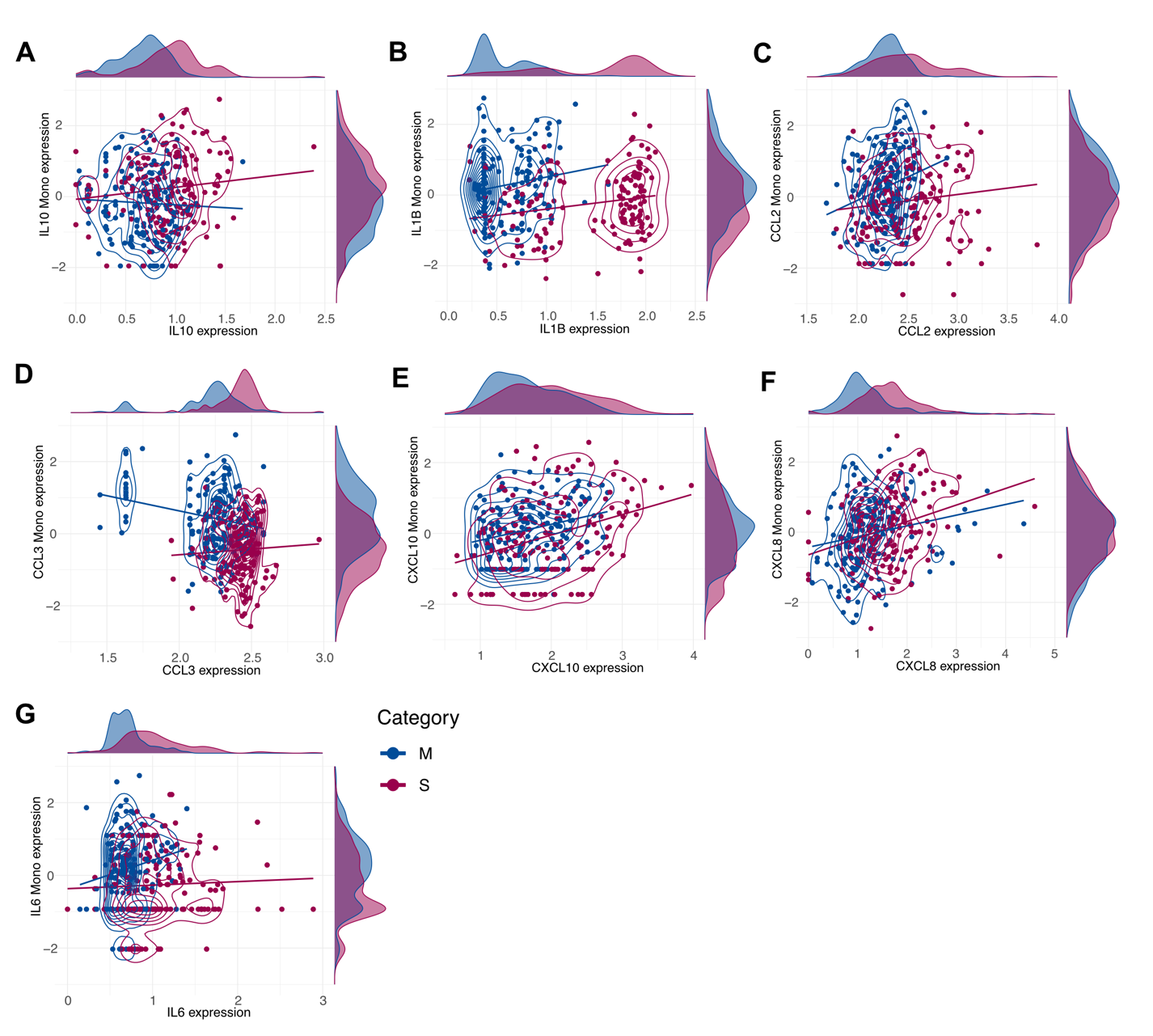

**Supplementary Figure 10.** Scatter plots of cytokines related to the COVID-19 pathway (ID: WP4891) with their RNA expression levels.

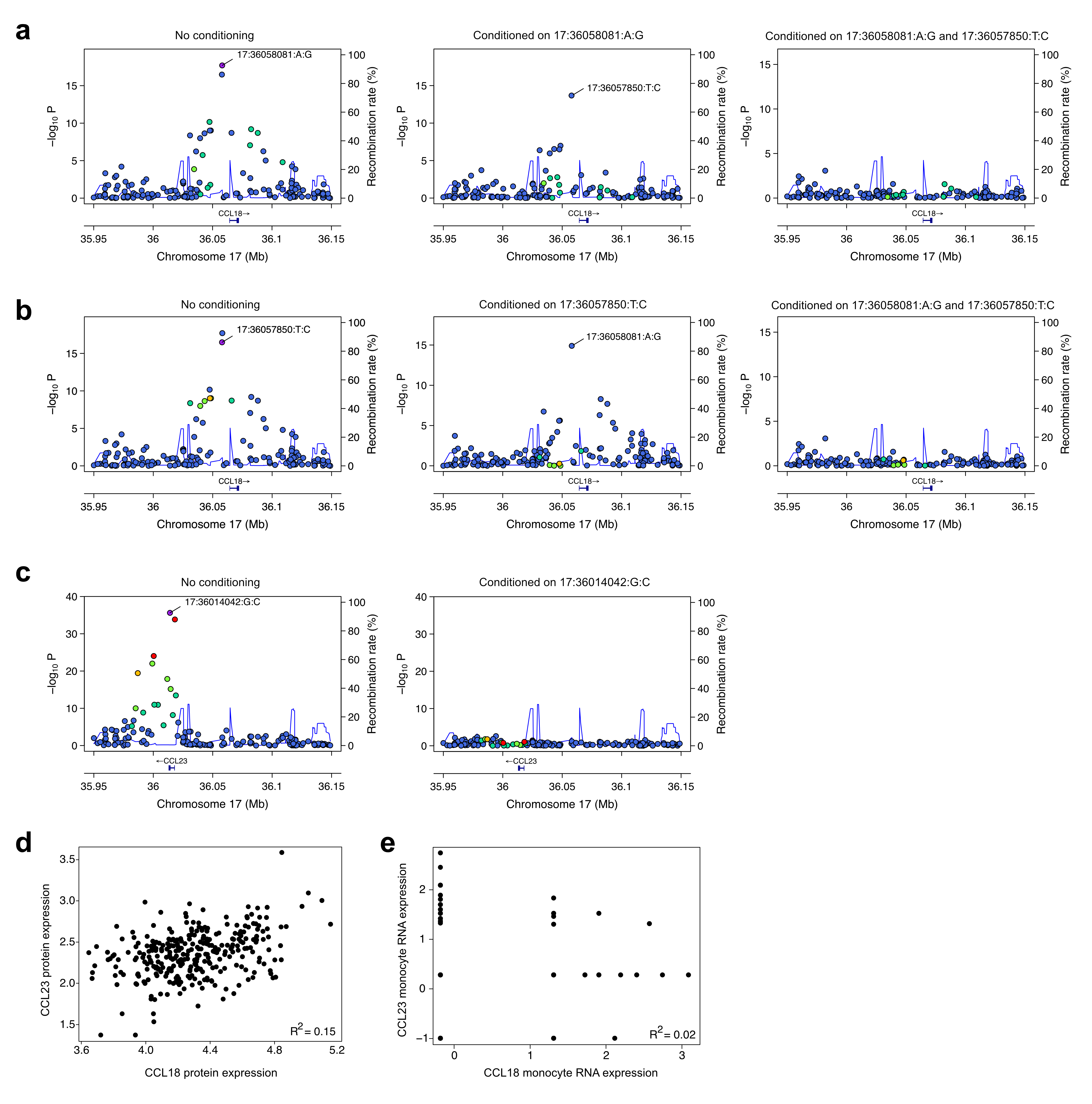

**Supplementary Figure 11.** Conditional analysis on pQTLs for CCL23 and CCL18. (a-c) Conditional analysis on CCL18 (a-b) and CCL23 (c). (d) Correlation between CCL18 and CCL23 protein expression levels. (e) Correlation between CCL18 and CCL23 RNA expression levels.

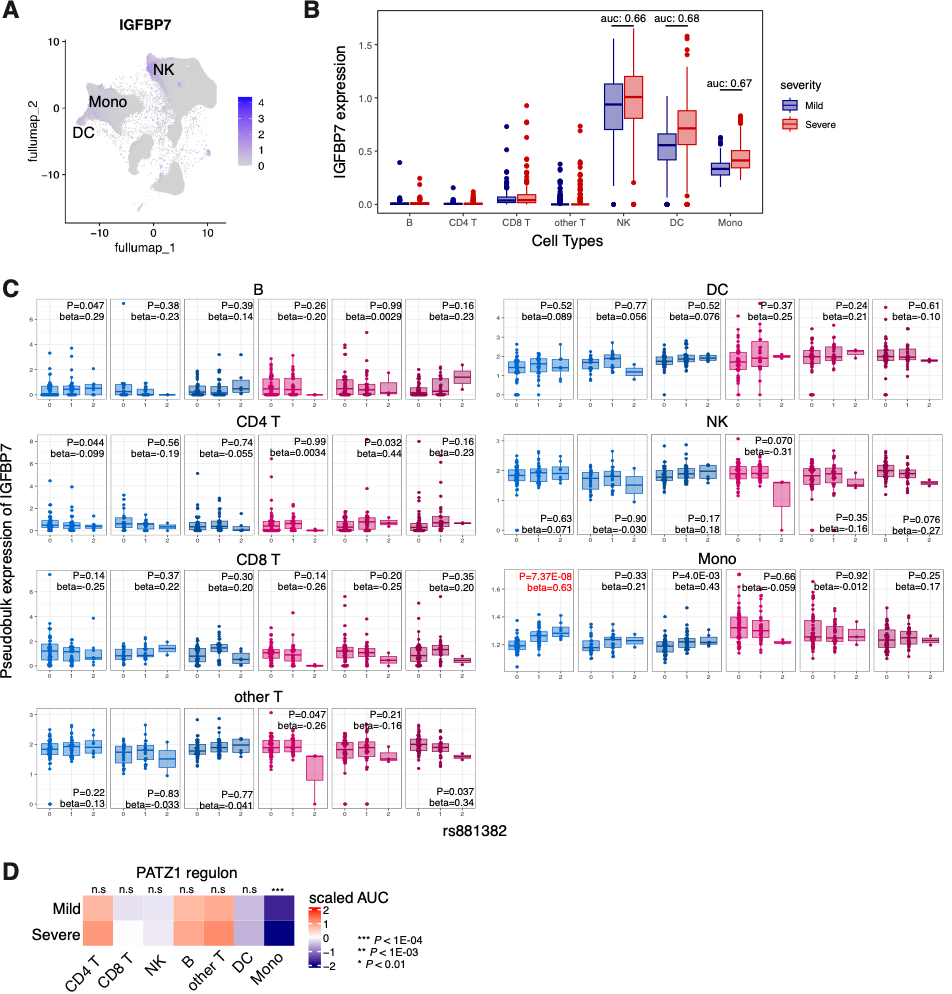

**Supplementary Figure 12.** Monocyte-specific regulation of *IGFBP7* expression mediating PATZ1 regulons. (A) *IGFBP7* expression projected on UMAP. (B) Differential expression of *IGFBP7* between the Mild and Severe groups across the cell types. Differential expression amount was calculated with the Presto package and annotated through the auc value. (C) TensorQTL summary statistics on the pair of *IGFBP7* and rs881382 across the cell types and conditions. (D) Scaled activity of PATZ1 regulon across the cell types and severity. Differential activity between severity was calculated by *t*-test for each cell type.

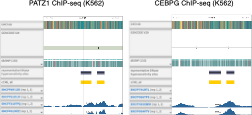

**Supplementary Figure 13.** ChIP-seq data for PATZ1 in the IGFBP7 locus in the K562 cell line. The track data were obtained from the ENCODE Genome Browser (ENCSR549PVK). The vertical line indicates the position of rs881382. The grey and yellow boxes represent DNase hypersensitivity sites and candidate *cis*-regulatory elements (cCREs), respectively. The subsequent tracks display conservative irreproducible discovery rate (IDR) threshold peaks, IDR threshold peaks, fold change over control, and signal *P*-values.
